# Self-Assembled Chambered Cardiac Organoids for Modeling Cardiac Chamber Formation and Cardiotoxicity Assessment

**DOI:** 10.1101/2024.07.25.605044

**Authors:** Xinle Zou, Fanwen Wang, Huilin Zheng, Tianci Kong, Duanqing Pei

## Abstract

The human heart, crucial for health and longevity, is a primary focus of medical research. Despite advancements in cardiac organoid technology, replicating early cardiac chamber formation stages remains challenging. Here, we develop chambered cardiac organoids (CCOs) by orchestrating cardiac signaling pathways, including the synergistic modulation of FGF and Wnt signaling via the HAND1 transcription factor. These CCOs exhibit stable chambers with self-organized outer myocardial and inner endocardial layers, and express developmental markers such as NKX2.5, TNNT2, and NAPPA, demonstrating physiological functionality including spontaneous contractions and calcium transients. Immunofluorescence and single-cell RNA sequencing confirmed the presence and stability of cardiomyocytes within CCOs. Further validation showed CCOs’ involvement in critical processes like endothelial-mesenchymal transition (EndoMT) and valvulogenesis. Ultrastructural and electrophysiological analysis revealed organized myofibrils and atrial-like action potentials. Importantly, CCOs proved effective in assessing cardiotoxicity through observable morphological changes, demonstrating specific cellular responses to established cardiotoxic compounds, thereby highlighting their potential for drug testing and safety evaluations. Our findings offer insights into cardiac chamber formation mechanisms and establish a robust platform for drug testing, disease modeling, and personalized medicine, representing a significant advancement in functional cardiac organoid development for both basic research and translational applications.

## Introduction

The human heart is uniquely structured and plays a vital role in mediating the circulation of not only nutrients but also the removal of wastes for an average of 80 years or so, a formidable challenge to model in animals ^1^. During heart development, the formation of cardiac chambers represents a critical stage where the initial tube-like structure of the heart differentiates into distinct atria and ventricles. This process is crucial for ensuring proper functional development of the heart through coordinated genetic pathways, signaling molecules, and cellular interactions ^2^. Initially, the heart consists of the endocardium and myocardium layers during tube formation and looping. Subsequently, the epicardium migrates from the proepicardial organ to envelop the developing heart ^3^. To model this developmental stage, cardiac organoids should exhibit at least four essential characteristics: stable chambers, well-defined inner endocardium and outer myocardium layers, expression of specific developmental markers, and demonstration of physiological functional characterization.

Over the past two decades, the emergent paradigm of using pluripotent stem cells (PSCs) to generate mini-organ/tissue or in vitro copies of human tissues and organs has provided much-needed progress, especially in modeling various aspects of the human heart^4–6^. Indeed, several well-established models have provided much-needed insights into organogenesis and pathology in vitro ^7–12^. Pertinent to the human heart, self-assembled chambered cardiac organoids (CCOs) have been reported ^7–10^. Especially Mendjan and colleagues reported the first 3D human chambered cardiac organoids, known as “cardioids” ^9^. Subsequently, specialized protocols for chambered outflow tract (OFT), chambered atrial, and chambered ventricular organoids, as well as other robust chambered cardiac organoids, have been developed ^13,14^. However, these CCOs often lack stable chamber, distinct outer myocardium and inner endocardium layers, or self-organization. Consequently, there remains a need for an ideal cardiac organoid model that can accurately represent early cardiac chamber formation stages.

Here, we addressed this construction challenge by orchestrating cardiac signaling pathways, including synergizing fibroblast growth factors (FGFs) ^9,13–17^, a well-known cardiac signal factor, with Wnt signaling, facilitated through the HAND1 transcription factor, to successfully construct the CCOs. These optimized CCOs successfully meet all four aforementioned criteria by accurately modeling heart development and also providing a robust platform for testing drug toxicity.

## Results

### Chambered cardiac organoids (CCOs) exhibit stable chamber structure

As mentioned in introduction, to accurately model the stage of heart chamber formation, cardiac organoids should at least exhibit four essential characteristics: stable chamber structure, well-defined inner endocardium and outer myocardium layers, expression of specific developmental markers, and demonstration of physiological functionality.

While heart chamber formation is reported to depend on the Wnt-BMP signaling axis, it is not sufficient to generate stable chambers with well-defined inner endocardium and outer myocardium layers ^9,13,14^, which is consistent with our own research (Fig.1a). Studies using model organisms and in vitro models have outlined the signaling pathways necessary for cardiomyocyte generation, including activin A (AA), retinoic acid (RA), BMP4, Wnt, FGF, TGFβ, among others^17–19^. In our effort to model ideal cardiac organoids at the heart chamber formation stage, we systematically screened those cardiac signals, across various stages of heart development-from pluripotency and mesoderm to cardiac mesoderm, cardiomyocyte formation, chamber formation, to chamber maintenance (Fig. 1b and Extended Data Fig. 1a). As a result, we successfully established ideal chambered cardiac organoids to effectively model this developmental stage.

**Figure 1.**
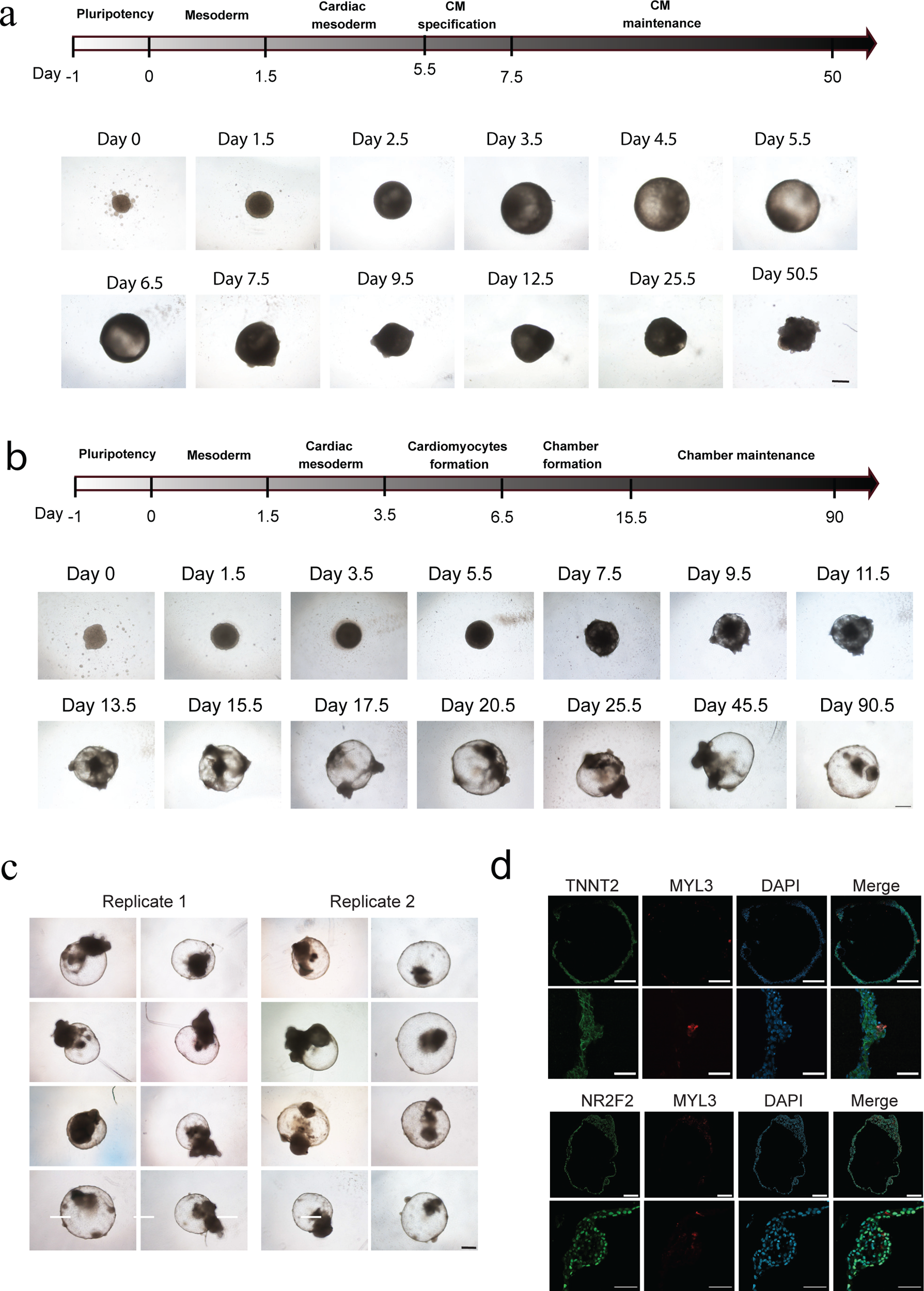
Stable chamber structure of cardiac organoids (CCOs) derived from hPSCs. a. Protocol and time-lapse of unstable chambered cardiac organoids formation captured in live brightfield images using Hofbauer’s protocol^9^. Scale bar, 500 μm. b. Protocol and time-lapse of CCOs formation captured in images using our protocol. Scale bars, 500 μm c. Representative brightfield images on day 90.5 from two independent biological replicates. Scale bar, 500 μm. d. Cryosections of CCOs on day 56.5, displaying cavity formation alongside the expression of cardiomyocyte marker TNNT2, ventricular marker MYL3, and atrial marker NR2F2. Scale bar, 200 μm.

We initial confirmed the formation and stability of chambers during CCOs formation through naked-eye observation under bright-field microscopy. Our results demonstrate that CCOs consistently exhibit stable chamber formation in 100% of organoids, with a single chamber appearing by day 11.5 and persisting beyond day 90.5 (Fig. 1b and 1c). Subsequently, immunofluorescence analysis revealed that these stable CCOs prominently feature *TNNT2* positive cells, indicative of cardiomyocytes, with *NR2F2* positive cells marking the atrial sector and a subset expressing a ventricular marker (Fig. 1d).

Furthermore, scRNA-seq analysis showed that 87.4% of cells within CCOs are cardiomyocytes (*TNNT2^+^*), among which 76.2% express the atrial marker *NR2F2* (Extended Data Fig. 1b). Throughout CCOs formation, a developmental progression was observed starting from pluripotency (marked by *NANOG^+^*, *POU5F1^+^*, and *SOX2^+^*)^20^, through mesoderm (marked by *TBXT^+^*, *MESP1^+^*, and *MIXL1^+^*)^21–23^, and cardiac mesoderm stages (marked by *HAND1^+^*)^24^, culminating in the stage of formation of cardiac chambers (Extended Data Fig. 1c).

Additionally, these CCOs transitioned into the first heart field (FHF) lineage (marked by *HAND1^+^*, *TBX5^+^*, *NKX2-5^+^*, and *TBX1^-^*)^25^ by day 6.5, preceding chamber formation (Fig. 1b and Extended Data Fig. 1c). Lastly, we validated our protocol in both the H1 and UiPS cell lines^20,26^, obtaining very similar results (Extended Data Fig. 1h, i), demonstrating the generalizability and applicability of our approach.

### CCOs exhibit distinct inner endocardium and outer myocardium layers and specific markers

Initially, the heart forms as a tube with well-defined inner endocardium and outer myocardium layers during looping, followed by epicardial migration from the proepicardial organ to cover the developing heart^3^. The endocardium plays a critical role in maintaining chamber stability and integrity during development and into adulthood ^27–29^. Thus, the presence of distinct outer myocardium and inner endocardium layers is pivotal during the heart’s chamber formation stage. However, previously reported cardiac organoids have not self-organized these layers, even with VEGF^9^.

Bulk RNA sequencing showed significant upregulation of endocardial markers such as CDH5^28^, PECAM1^28^, CD34^30^ and ESAM^8^, from day 3.5 in our CCOs (Figure 2A). Immunofluorescence analysis of cryosections further confirmed the architecture of outer myocardial and inner endocardial layers in our CCOs, identified respectively by TNNT2 and CDH5 (Fig. 2b). To evaluate the expression of specific developmental markers linked to chamber formation and heart looping during heart development, such as NPPA, NPPB, TBX5, HAND1 and HAND2 ^9,31–33^, we conducted a comprehensive analysis of time-course bulk RNA sequencing data. Our findings revealed significant upregulation of these marker genes across the formation stages of CCOs, underscoring their pivotal role in cardiac organoid development (Fig. 2c).

**Figure 2.**
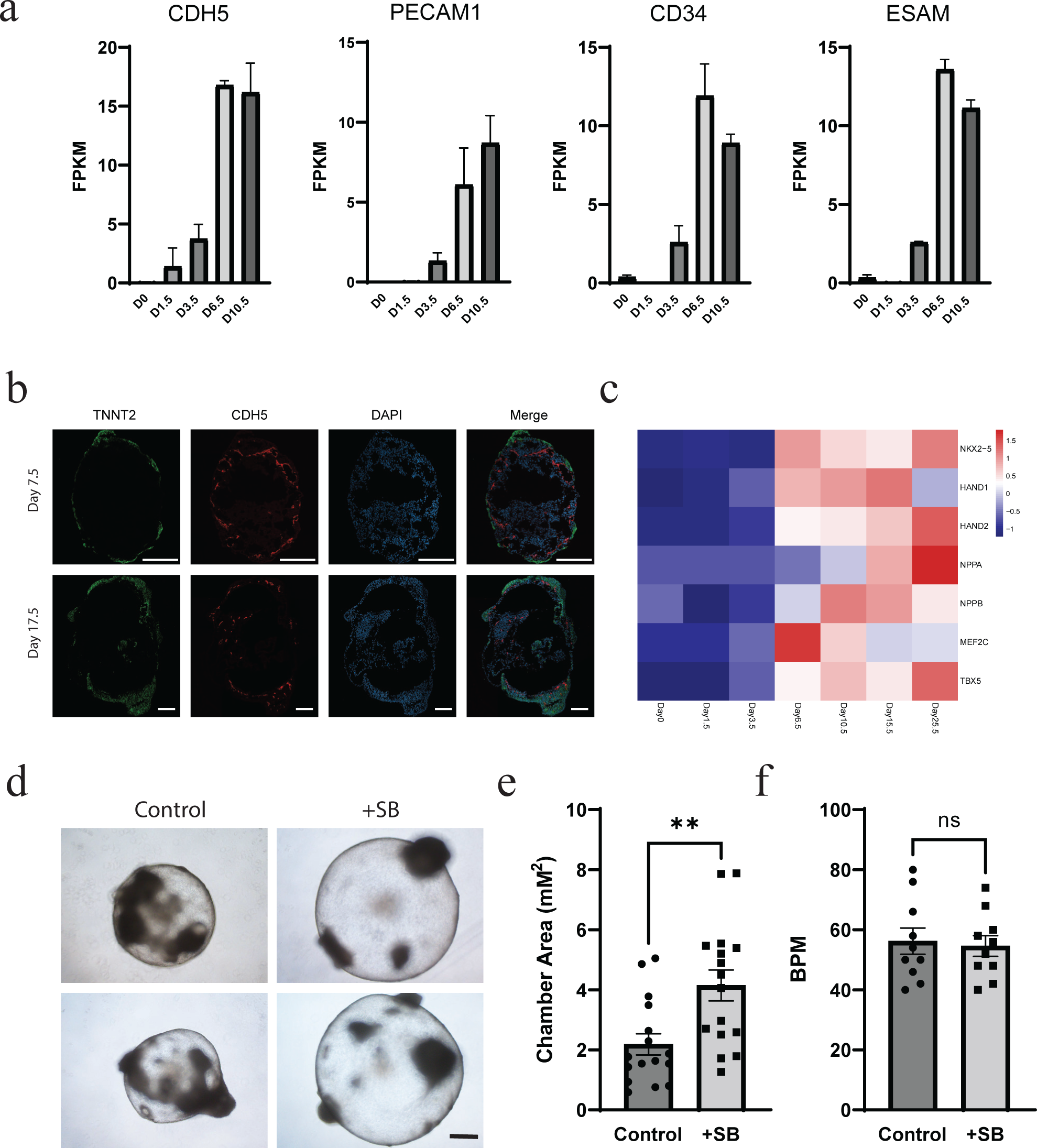
Inner endocardium and outer myocardium layers and markers in the CCOs. a. Expression levels of the endocardial specific markers during CCOs formation. FPKM, Fragments Per Kilobase of transcript per Million mapped reads. b. Example images illustrating the architecture of outer myocardial and inner endocardial layers in CCOs on day 7.5 and day17.5. Scale bar, 200 μm. c. Heatmap displaying gene expression of markers associated with chamber formation and heart looping during CCOs development from day 0 to day 25. d. Representative brightfield images of CCOs treated with or without 0.5 μM SB431542 for 20 days, captured on day 48.5. Scale bar, 500 μm e. Measurement of chamber areas in cardiac organoids on days 48.5 after treatment with 0.5 μM SB431542. f. Beating rate of CCOs treated with or without SB431542 on day 48.5. BMP, beats per minute. All bar graphs show mean ± SEM.

The endocardium also plays crucial roles in heart development and cardiovascular diseases through endothelial-to-mesenchymal transition (EndoMT), induced by transforming growth factor (TGF-β) ^34,35^. This transition is critical in the formation of key structures like the endocardial cushions, and these cushions ultimately develop into the valves of the heart, playing a critical role in preventing blood backflow and regulating the direction of blood flow within the heart ^36,37^. During the formation of cardiac conduction systems (CCOs), we have observed and verified an endothelial-to-mesenchymal transition (EndoMT) process. This transition is evidenced by the expression of specific markers such as *TWIST1*, *ACTA2*, *SANL2*, and *CDH2*, concurrent with the emergence of endocardial tissue identified by *CDH5* and *PECAM* (Fig. 2a) ^38^. To further confirm valve formation, we utilized COL1A1, a specific marker for heart valves. Immunofluorescence analysis revealed COL1A1-positive cells within the inner CCOs region (Extended Data Fig. 2b).

To investigate the functional role of endocardium in chamber formation, we treated CCOs with SB431542 (SB), a potent and selective inhibitor of the TGF-β signaling pathway, from day 28.5 to day 48.5. Remarkably, the SB-treated group exhibited chamber areas nearly twice as large as the control group (Fig. 2d, e, and Extended Data Fig. 2c), consistent with the known role of endocardium in maintaining the stability and integrity of the heart’s chambers ^27–29^. Interestingly, the beating rate was not affected by SB (Fig. 2f).

Finally, to simulate the process of epicardium immigration covering the naked developing heart ^3^, we generated epicardium from H9-tdTomato cell lines, randomly integrating tdTomato into the genome (Extended Data Fig. 2d). Remarkably, after aggregating epicardial cells using AggreWell400 plates, we co-cultured these aggregations with our developing heart model. As a result, we successfully generated CCOs encompassing epicardial, myocardial, and endocardial layers (Extended Data Fig. 2e, 2f).

Together, our cardiac conduction system model demonstrates a functional inner endocardial layer and expresses specific markers relevant to heart development stages, highlighting its potential for studying complex cellular interactions in cardiac development.

### CCOs exhibit functional physiological characterization

While our CCOs exhibit stable chamber formation, well-defined inner endocardium, outer myocardium structure, and express specific markers, they also require functional characterization. So, we conducted multi-level studies to assess the functional characterization of our CCOs. At the ultrastructural level, we show that these CCOs display hallmark features of cardiomyocytes, including well-organized sarcomeres, Z-lines, and interconnectedness via intercalated discs (ID) through immunofluorescence and electron micrographs (Extended Data Fig. 3a, 3b). At the single-cell level, with whole-cell patch clamp experiments, we demonstrate that individual cardiomyocytes possess atrial-like action potential profiles with typical features like action shape, resting membrane potential (RMP), action potential amplitude (APA), and action potential duration (APD) (Extended Data Fig. 3c, 3d).

Consistent with properties described above, we show that these CCOs have heart rates around 60 beats per minute (BPM) from day 21.5 to day 68 (Extended Data Fig. 3e, 3f), compatible for future therapeutic development, as synchronizing the beating rates of transplanted organoids with the patient’s heart rhythm is essential for effective integration and function within the recipient’s cardiac system. Additionally, these CCOs also exhibit robust beating strength, as demonstrated by visible and powerful contractions observed in videos and calcium imaging (Extended Data Fig. 3f, and Videos 1, 2, and 3).

In summary, CCOs have been successfully developed to model critical chamber formation stages of heart development, demonstrating stable chamber formation with distinct inner endocardium and outer myocardium layers. These organoids also exhibit functional physiological characteristics akin to human cardiomyocytes, making them a robust platform for studying cardiac development and diseases.

### Cellular composition analysis of CCOs using single-cell sequencing

To determine cell composition of our CCOs, we performed single-cell sequencing analysis on both 2D and 3D cultures on day 43.5. We obtained 25,274 and 14,614 cells from 2D and 3D cultures after stringent quality control respectively. Afterwards, normalization, batch-effect correction, dimensionality reduction, and clustering were performed, and cell annotation was performed based on the marker gene. Specifically, there are five cell types in 2D cultures: atrial cardiomyocytes, ventricular cardiomyocytes, cardiac progenitors, valve and fibroblasts (Fig. 3a, 3b and Extended Data Fig. 4a). Interestingly, two additional cell types, endocardium expressing *CDH5*, and epithelial progenitors expressing *GABRP* and *GRHL2*, are present in the 3D cultures (Fig. 3a-3c and Extended Data Fig. 4a-d).

**Figure 3.**
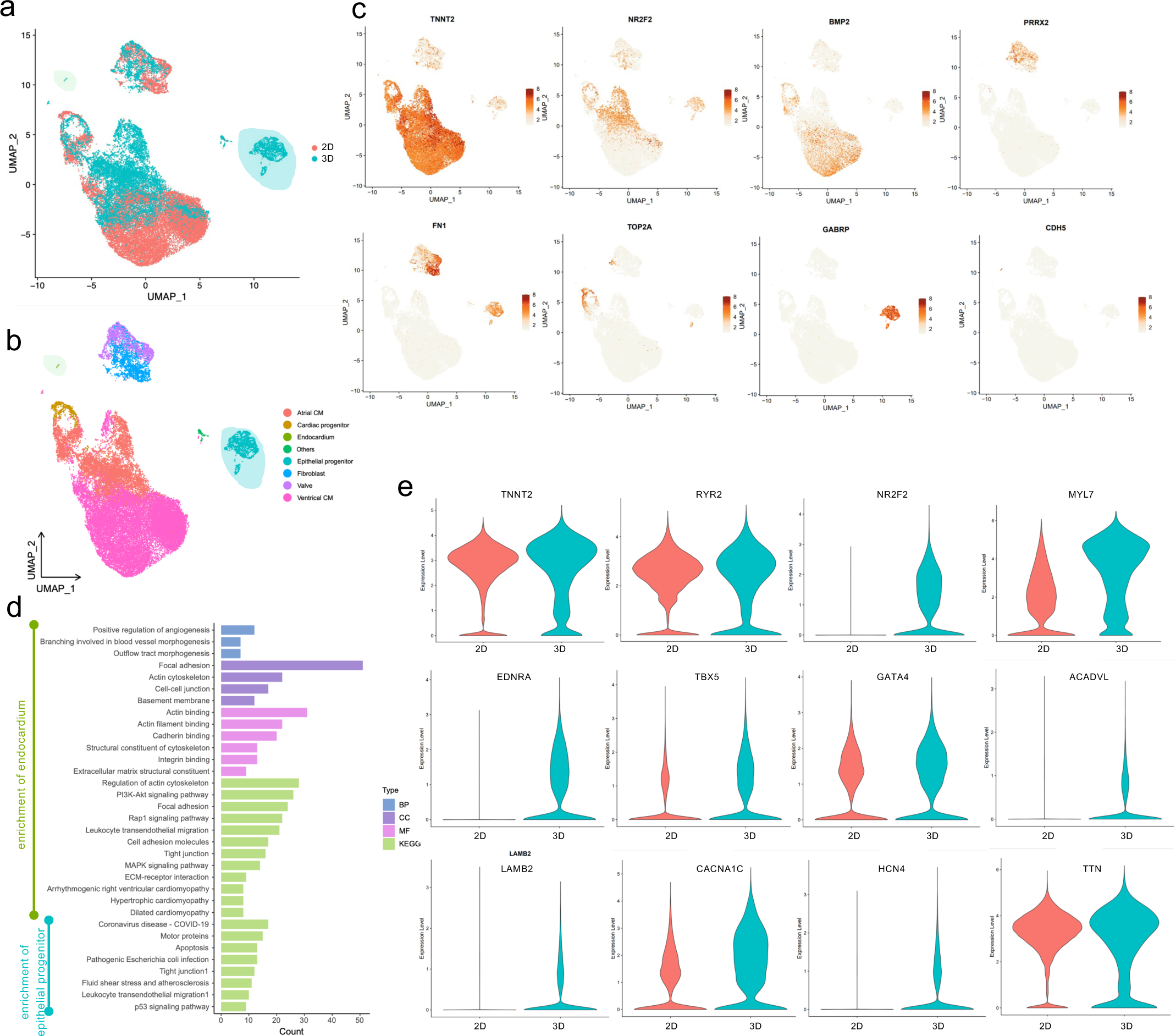
Comparative Analysis of Cellular Diversity and Gene Expression Profiles in 2D and 3D Cell Cultures. a. Data Distribution: Illustrates the distribution of cells in 2D and 3D culture samples. Cells in 2D samples are denoted in red, while those in 3D samples are depicted in blue, facilitating visualization of cell distribution within each sample in the dataset. b. Cell Type Annotation Visualization: Offers a UMAP visualization of cell type annotation using single-cell RNA-seq data from both 2D and 3D cultures, with colors representing different cell types. c. Marker Gene Expression: Presents a UMAP scatter heatmap depicting marker genes for various cell types, with color gradients indicating expression levels. Cellular annotation was conducted based on marker gene expression in different clusters. d. Enrichment analysis: Based on annotation results, 3D cultures demonstrate a greater diversity of cell types compared to the 2D group. Differential genes specific to endocardial and epithelial progenitor cells in 3D cultures were enriched and analyzed. e. Gene Expression Analysis: Compares the differential expression of genes marking cardiac maturation between the 2D and 3D culture groups, alongside violin plots illustrating the distribution range of expression levels for key genes.

The endocardium and epithelial progenitor cells are of particular interest as they are specific to the 3D cardioids. Based on GO and KEGG analysis of the highly variable genes, we show that the endocardial cells are implicated in angiogenesis, outflow tract morphogenesis, focal adhesion, and related signaling pathways such as Rap1, MAPK, and PI3K-akt. Conversely, the epithelial progenitors are associated with processes such as apoptosis, motor proteins, tight junction regulation, and the p53 signaling pathway (Fig. 3d). Comparison of gene expression between 2D and 3D cultures uncovers significant expression of relevant genes such as ion channels (*RYR2*, *CACBA1C*, *HCN4*), progenitor (*EDNRA*), β-oxidation (*ACADVL*), as well as genes associated with cardiomyocytes (*TNNT2*, *MYL7*, *NR2F2*, *TBX5*, *GATA4*, *LAMB2*, *TTN*) specific to 3D culture (Fig. 3e).

Additionally, both atrial and ventricular cardiomyocyte types are present in both 2D and 3D cultures. Further enrichment analysis of highly expressed genes in 3D versus 2D cultures reveals their involvement in Z disc organization, heart rate regulation, and heart development, along with associated signaling pathways such as MAPK and PI3K-Akt (Extended Data Fig. 5e, 5f). Furthermore, comparing the expression differences of maturation marker genes in atrial or ventricular cells between 2D and 3D cultures demonstrated significant increases in ion channels, progenitor cells, and β-oxidation processes in the 3D cultures (Extended Data Fig. 5g, 5h).

In conclusion, these results confirm the broader cell diversity and functional maturation generated by the 3D culture system.

### FGFs play critical roles in the formation and stability of chambers in CCOs

Chambers are significant features during the stage of heart chamber formation in heart development. Although FGFs have been shown to play crucial roles in heart development^15,16^, how this crucial signaling system impacts chamber formation remains unknown. To investigate its role, we tested the synergistic effects of FGF and Wnt pathway inhibitors in our system. We specifically activated the FGF-Wnt pathways with FGF2 and CHIR99021, and inhibited these pathways using PD173074 (PD) and IWP2. Given the high expression of FGFR1 (Extended Data Fig. 5d), we initially tested the FGFR1 inhibitor PD173074 ^39^. Our data suggest that inhibiting FGFR1 enables the formation of CCOs with distinct chamber structures (Fig. 1b, 1c, 4a-4c and Extended Data Fig. 5a). Without PD, the organoids exhibit no chamber formation, as shown in the brightfield images (Fig. 4a, 4b). Dynamic imaging confirms the lack of chamber formation over time (Extended Data Fig. 5a). Consistently, we show with immunofluorescence analysis that the organoids are TNNT2 positive, despite diminished chamber sizes without PD (Fig. 4c). We also tested PD166866, a selective inhibitor of FGFR1 tyrosine kinase activity, and H3B-6527, a highly selective covalent inhibitor of FGFR4, and show that PD166866 behaves similarly as PD173074, while H3B-6527 does not, suggesting that FGFR1 tyrosine kinase activity, rather than FGFR4, is critical for chamber formation (Extended Data Fig. 5b, 5c). The reason may be differences in the expression levels of FGFRs (Extended Data Fig. 5d).

**Figure 4.**
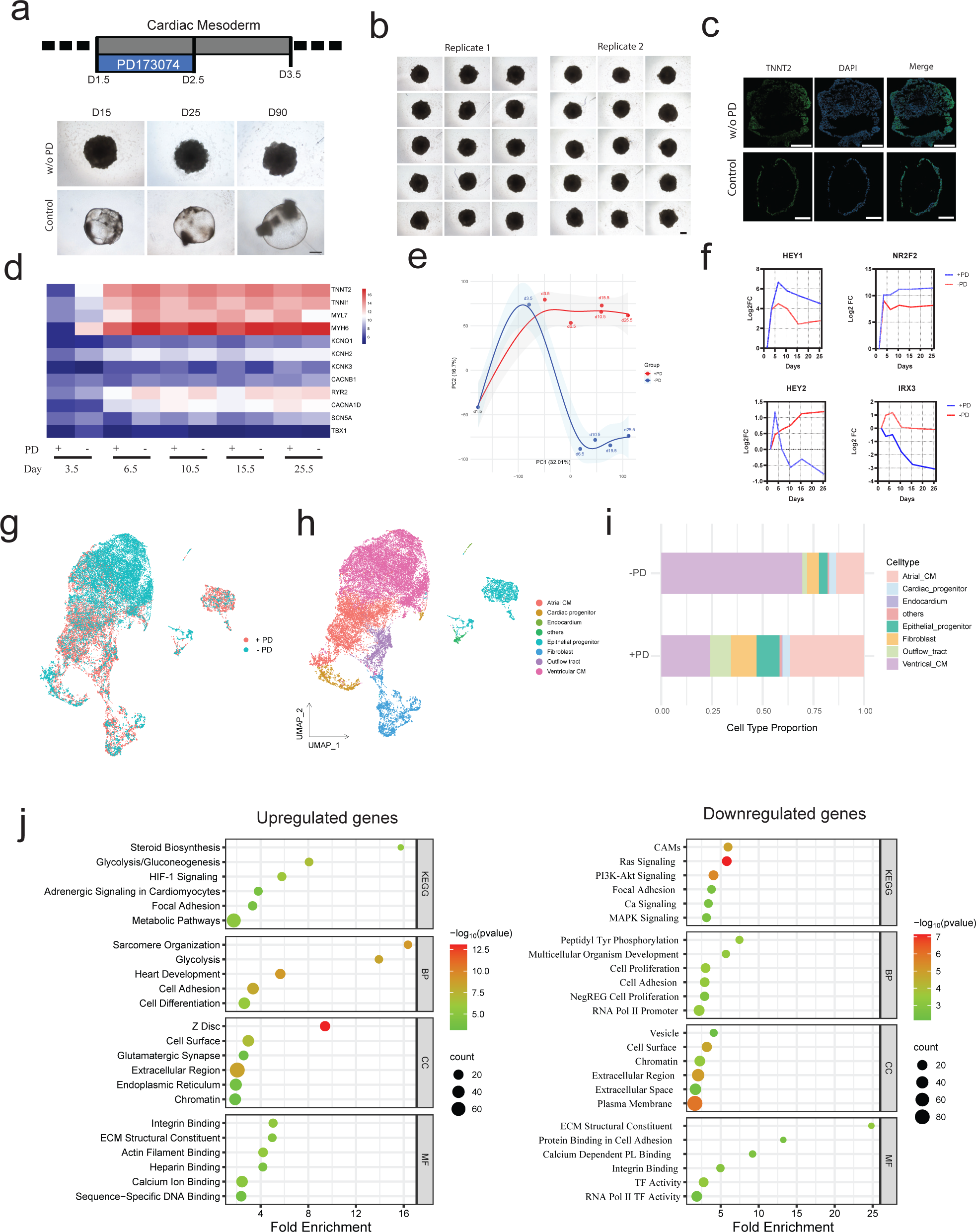
Inhibition of FGF is crucial for CCOs formation during cardiac mesoderm induction stage. a. Chamber formation of CCOs is dependent on PD173074 (PD). Scale bar, 500 μm. b. Brightfield images on day 16.5, showing solid cardiac organoids structures across three independent biological replicates without PD. Scale bar, 500 μm. c. Cryosections of CCOs on day 16.5 (with PD and without PD). Scale bar: 200 μm. d. Heatmap illustrating the expression patterns of key genes, including cardiomyocyte markers and ion channels, during cardiomyocyte differentiation with and without PD. VST, variance-stabilized transformed counts. e. PCA plot illustrating distinct cell fates with and without PD. f. Cardiac-specific cell type gene expression analysis (log2 fold-change vs. day 1.5) in CCOs. g. Data Distribution: Illustrates the cell distribution of CCOs with or without PD. Cells with PD are denoted in red, while those samples without PD are depicted in blue, facilitating visualization of cell distribution within each sample in the dataset. h. Cell Type Annotation Visualization: Offers a UMAP visualization of cell type annotation using single-cell RNA-seq data from cultures, with colors representing different cell types. i. Horizontal bar charts depicting the cell proportion of each cell cluster with or without PD groups. j. GO and KEGG terms upregulated or downregulated in CCOs with PD as control on day 3.5.

A heatmap of time course data demonstrates that cardiomyocyte differentiation with or without PD is not affected (Fig. 4d). On the other hand, PCA analysis reveals a distinct alteration in organoids cell fate induced by PD (Fig. 4e). By volcano plot analysis, we show that key genes associated with cardiomyocyte differentiation are significant upregulated at the cardiac mesoderm stage on day 3.5, particularly in samples without PD treatment (Extended Data Fig. 5e). However, by day 10.5 during chamber formation, we observe a shift in cardiomyocyte fate: increased expression of atrial-specific genes like NR2F2 in organoids without PD, and elevated levels of ventricular-specific genes, notably IRX4 in those with PD (Extended Data Fig. 5e), confirmed by more detailed cardiac-specific cell type gene expression analysis (Fig. 4f). Additionally, single-cell sequencing analysis indicates a notable change in the proportion of cell clusters. Particularly, the addition of PD results in increased levels of endocardium, fibroblasts, outflow tract cells, epithelial progenitors, and atrial cardiomyocytes (Fig. 4g-4i, and Extended Data Fig. 5f). These data are consistent with previous reports that FGF signaling acts upstream of NKX factors to maintain ventricular identity^40^.

Gene Ontology (GO) and Kyoto Encyclopedia of Genes and Genomes (KEGG) enrichment analyses on day 3.5 revealed significant enrichment in various biological processes, cellular components, and molecular functions for both upregulated and downregulated genes. Upregulated genes are enriched in processes like cholesterol biosynthesis, heart development, and glycolysis, as well as metabolic pathways and adrenergic signaling, while downregulated genes associated with processes such as cell proliferation and adhesion, and pathways like Ras signaling and PI3K-Akt signaling. The downregulated genes are involved in cell proliferation and adhesion, and their proteins in cellular components like the plasma membrane and chromatin (Fig. 4j). These results provide insights into the regulatory mechanisms governed by FGFs leading to CCOs formation.

### HAND1 is indispensable for the formation of CCOs

As mentioned above, PD173074 is critical for the CCOs formation. To further investigate the mechanism of CCOs formation, we conducted comprehensive gene expression analyses across five time points. We tracked changes in sets of upregulated and downregulated transcription factors. Utilizing Venn diagrams, we observed dynamic shifts in transcription factor expression, with some genes exhibiting stability over time. We identified 9 upregulated transcription factors and 4 downregulated transcription factors (Fig. 5a). Additionally, we ranked the log2FC of 13 selected transcription factors in cardiac organoids without PD173074 as the control on day 3.5 (Fig. 5b and Extended Data Fig. 6a), considering their expression levels (Fig. 5c, 5d). Ultimately, we identified HAND1 and NKX2-5, two well-known transcription factors involved in cardiac development, as potential transcription factors for further research. Notably, it has been reported that HAND1 plays crucial roles in cardiac chamber formation^9,31,32^. To this end, we first generated two HAND1 knockout (KO) H9 cell lines labeled as #9 and #15, alongside two NKX2.5 knockout H9 cell lines labeled as #14 and #24. Brightfield images and immunofluorescence analysis of cryosections revealed that HAND1 KO cells exhibit solid organoids without discernible chambers, yet they remain TNNT2-positive. In contrast, NKX2.5 KO cells generate chambered structures (Fig. 5e, 5f, and Extended Data Fig. 6b, 6c). These findings strongly suggest that HAND1 is indispensable for the chamber formation of CCOs, while NKX2.5 is not. These results also suggest that heart chamber formation is independent of cardiomyocyte differentiation.

**Figure 5.**
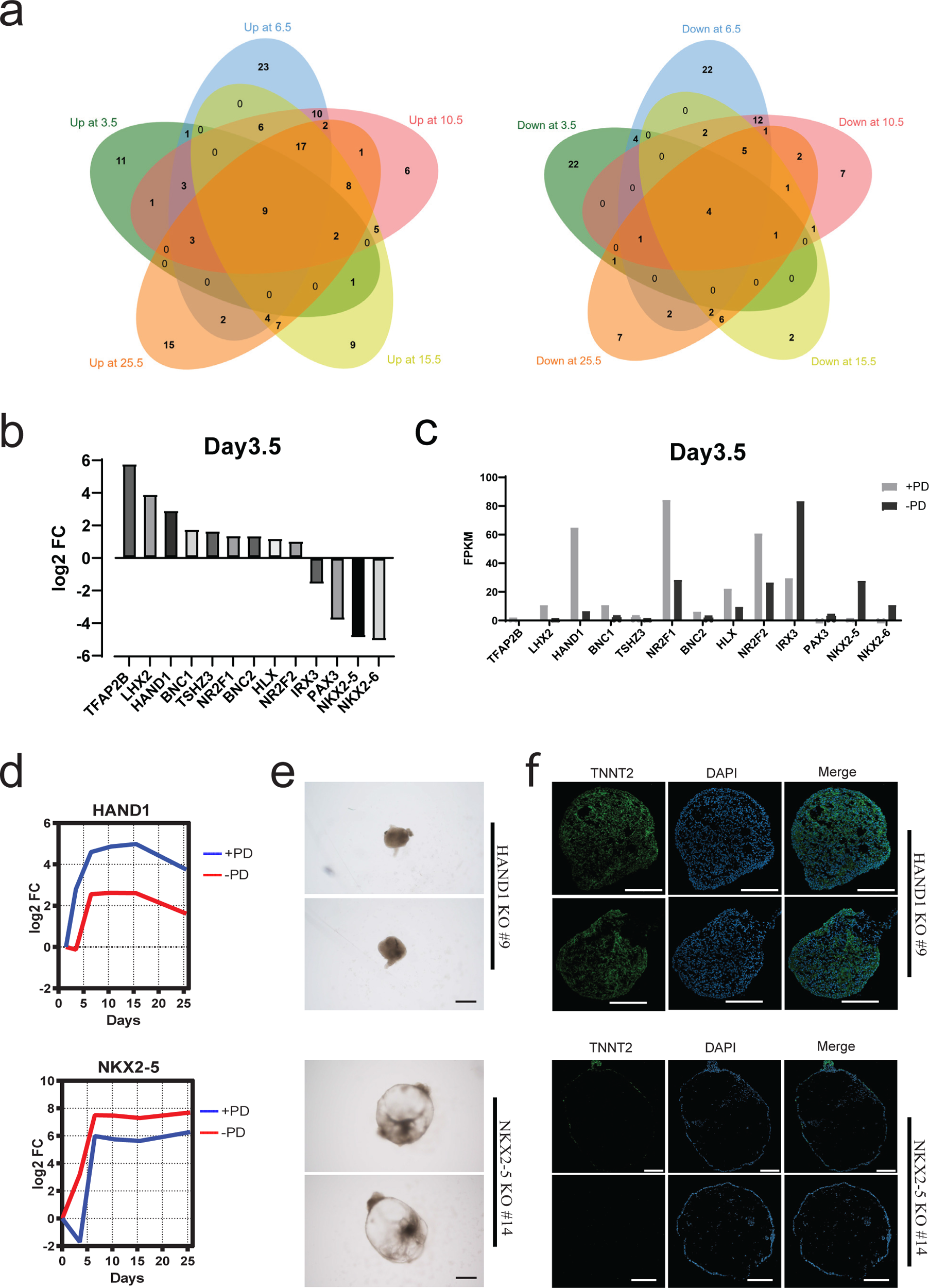
HAND1 is essential for the formation of CCOs. a. Venn diagram illustrating upregulated or downregulated transcription factor genes in cardiac organoids on different days, without PD173074 as control. b. Bar chart of log2 fold change of 13 selected transcription factors in cardiac organoids without PD173074 as the control on day 3.5. c. Expression levels of the selected transcription factors on day 3.5 of cardiac organoids. FPKM, Fragments Per Kilobase of transcript per Million mapped reads. d. HAND1 and NKX2.5 gene expression analysis (log2 fold-change vs. day 1.5) with or without PD173074 in cardiac organoids. e. Representative brightfield images of HAND1 and NKX2.5 KO cardiac organoids. Scale bar, 500 μm. f. Confocal cryosection images of HAND1 and NKX2.5 KO cardiac organoids stained with TNNT2. Scale bar, 200 μm.

### AA and RA are essential for chamber formation or stability in CCOs

AA and RA are both critical signaling molecules in cardiac development, including the induction of mesoderm and cardiac mesoderm^17,41,42^. However, their specific roles in chamber formation and stability remain unclear. After systematically testing AA and RA at various concentrations (A4R50: AA 4 nM and RA 50 nM, A4R500: AA 4 nM and RA 500 nM, A50R500: AA 50 nM and RA 500 nM), we observed that all three groups induced chamber formation in the CCOs (Fig. 6a). However, in the low AA concentration groups (A4R50 and A4R500), the chambers remain stable regardless of varying levels of RA. Notably, the A4R500 group exhibited a larger chamber compared to the A4R50 group (Fig. 6c), suggesting that higher RA concentrations facilitate chamber enlargement.

**Figure 6.**
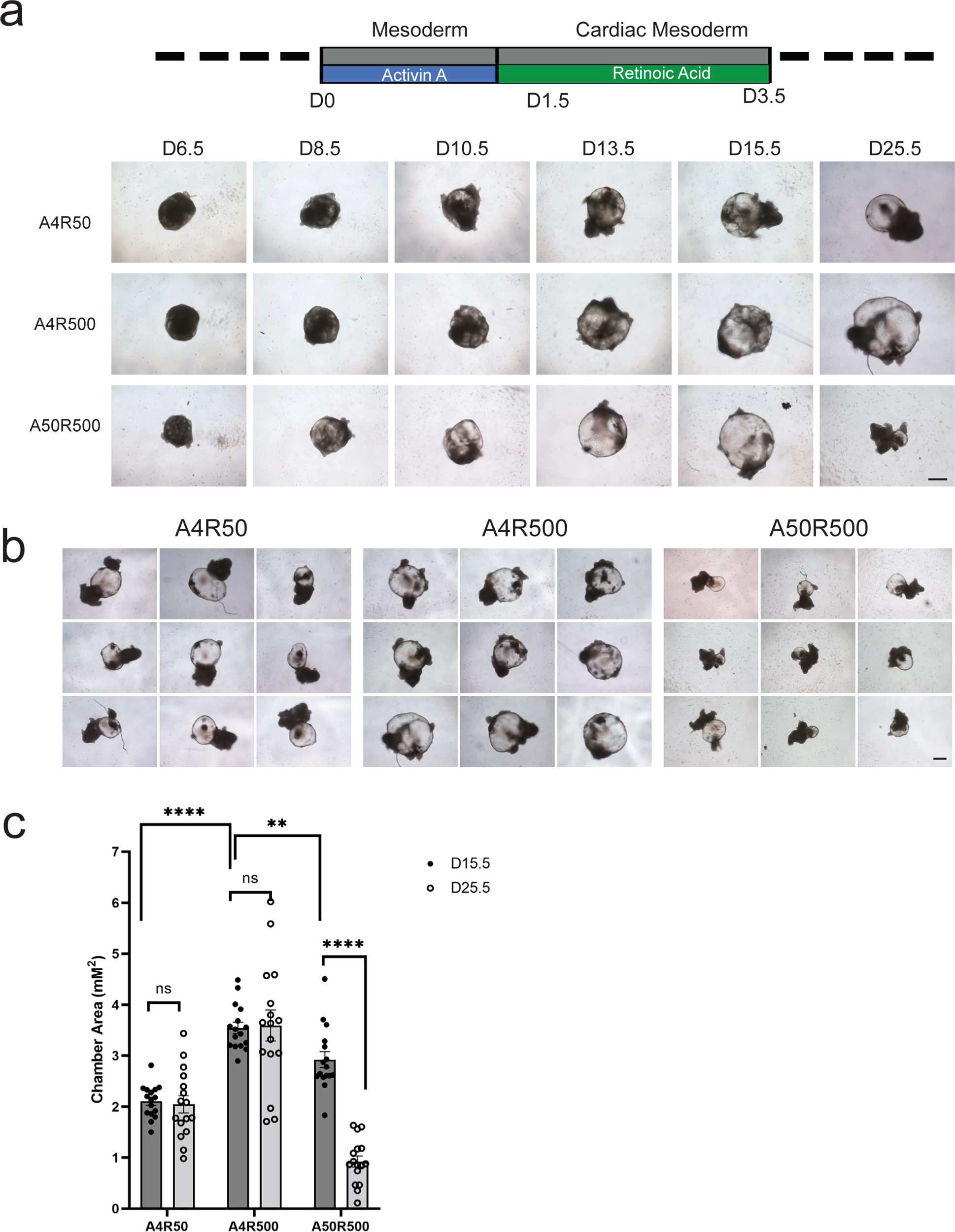
The roles of activin A (AA) and retinoic acid (RA) in the chamber formation and stability of CCOs. a. Time-lapse brightfield images showing the development of CCOs under different concentrations of AA and RA: A4R50 (AA 4 nM and RA 50 nM), A4R500 (AA 4 nM and RA 500 nM), and A50R500 (AA 50 nM and RA 500 nM). Scale bar, 500 μm. b. Representative brightfield images of CCOs induced with varying concentrations of AA and RA on day 25.5. Scale bar, 500 μm. c. Measurement of chamber areas in CCOs on days 15.5 and 25.5 after treatment with different concentrations of AA and RA.

Furthermore, when we increased the AA level from 4 nM to 50 nM (A4R500 to A50R500), the chamber areas of CCOs significantly decreased from day 15.5 to day 25.5, suggesting that lower AA concentration are conductive to chamber stability (Fig. 6). Given that AA can promote the efficiency of cardiomyocyte differentiation^41^, we adopted A10R500 as our protocol.

Together, these results that in addition to FGFs, both AA and RA are critical to chamber formation and stability in a dose dependent fashion.

### Cardiotoxicity assessment platform using human CCOs

Cardiac organoids with chamber have the potential to serve as a testing platform for drugs in terms of efficacy and toxicity^10,43^. Our CCOs, distinguished by their organized structure and extended maintenance capabilities, significantly enhance reliability and suitability for drug efficacy and toxicity assessments. This stable chambered design allows precise monitoring of chamber-specific morphological changes under brightfield microscopy images in response to treatments such as thalidomide and acitretin, emphasizing their utility in drug toxicity screening. Thalidomide is infamous for causing severe birth defects in newborns^44^, and acitretin, a known teratogens, significantly impact heart development and poses serious risks to fetal cardiac health^45,46^. Aspirin, with well-documented anti-inflammatory and pain-relieving properties, served as a negative control.

Our observations reveal that both thalidomide and acitretin induce severe morphological defects compared to Aspirin and wild type control (Fig. 7a and Extended Data Fig. 7a). By day 20.5, the treatment groups exposed to thalidomide and acitretin exhibit significantly smaller chamber areas compared to the control (Fig. 7a). Specifically, the thalidomide group displays clear chamber formation from day 6.5 to day 15.5, but fails to maintain this structure, transitioning to a spherical shape by day 20.5. Additionally, our findings suggest that thalidomide affects our CCOs during the mesoderm stages, while acitretin appears to impact the pluripotent stage (Fig. 7d). Conversely, the acitretin group exhibits an irregular shape as early as day 6.5 (Fig. 7a and Extended Data Fig. 7a). Moreover, both treatment groups exhibit either no or severely reduced beating rates and chamber area (Fig. 7b, 7c).

**Figure 7:**
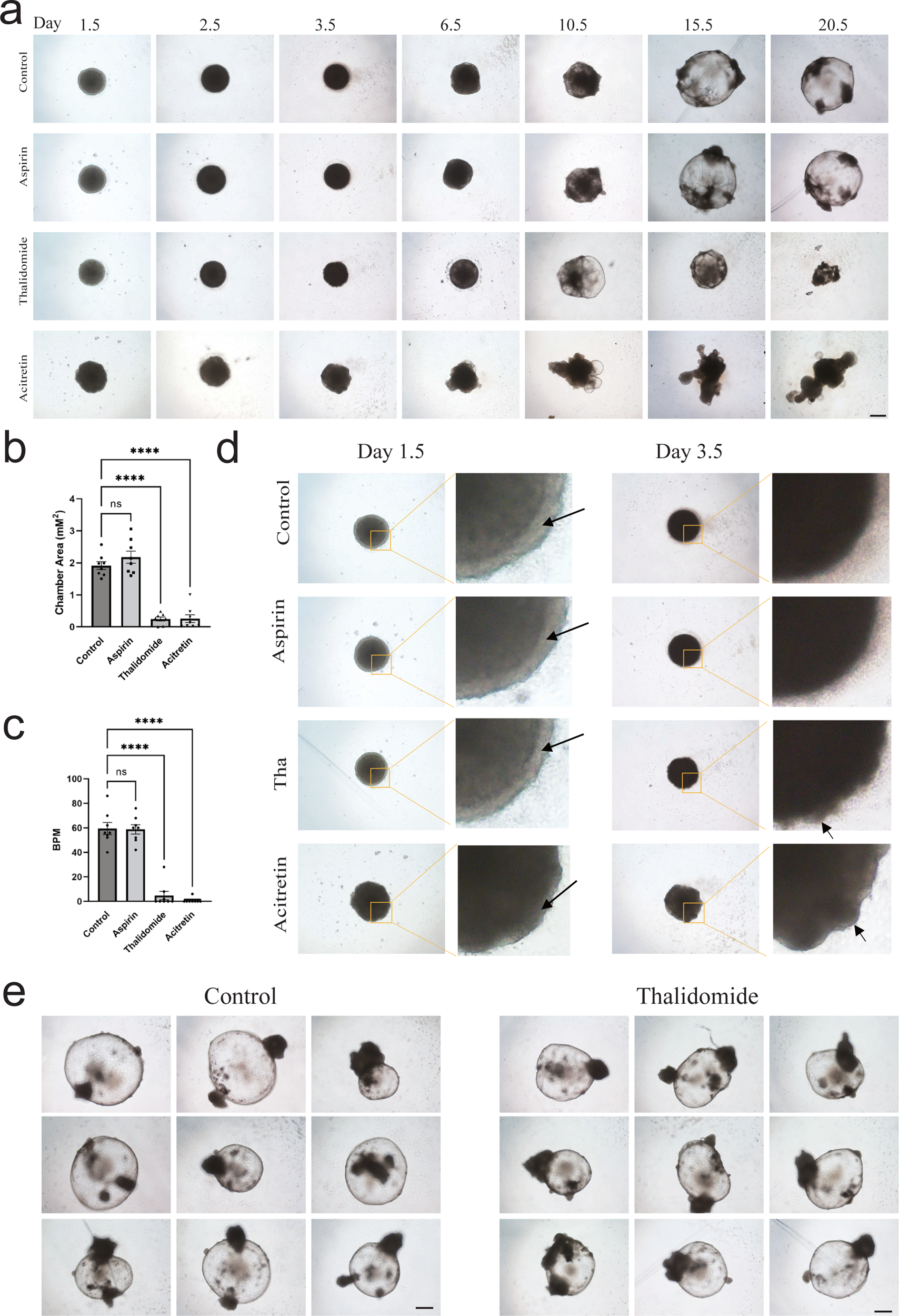
Establishment of CCOs platform for cardiotoxicity screening. a. Live brightfield images of CCOs treated with different drugs. All drugs were added into the CCOs from day 0. Aspirin, 30 μM; thalidomide, 10 μg/ml; acitretin, 50 nM. Scale bar, 500 μm. b. Quantification of the chamber areas of CCOs treated with or without drugs. c. Beats per minute of CCOs treated with or without drugs. d. Enlarged brightfield images of CCOs treated with different drugs on day 1.5 and day 3.5. e. Live brightfield images of CCOs captured on day 36.5, treated with thalidomide starting from day 15.5. Scale bar, 500 μm.

Thalidomide primarily induces embryonic damage during early developmental stages, typically between days 20 and 36 after fertilization^47^. To further investigate the timing window of thalidomide exposure, we administered thalidomide to the CCOs starting from day 15.5, once CCOs formation is complete, and continued the exposure until day 36.5. The results show that thalidomide does not alter the chamber area when applied after day 15.5 (Fig. 7e and Extended Data Fig. 7b), and the beating rates of the CCOs remain unaffected (Extended Data Fig. 7c). While our data suggest that thalidomide’s impact on CCOs on day 15.5 is not significant, it is important to note that late fetal exposure in animal models, such as rats, has been shown to induce brain damage^48^, indicating a potential lack of a safe time window for this drug. Together, we demonstrate that these CCOs may be further optimized for cardiotoxicity assessment.

## Discussion

We describe the creation of chambered cardiac organoids (CCOs) derived from human pluripotent stem cells (hPSCs), which simulate the stages of heart chamber formation by coordinating multiple cardiac signaling pathways, including synergistic FGF and Wnt pathways mediated by HAND1. These CCOs demonstrate robust responsiveness to drug treatments in toxicity screenings, underscoring their effectiveness in pharmacological testing. Further studies may extend this approach to generate cells, tissues, and miniature organs for developmental, pharmacological, and pathophysiological modeling.

The Wnt/β-catenin signaling pathway, along with RA and AA has been implicated in cardiac development at stages such as mesoderm specification, myocardial differentiation, and proliferation^41,42,49,50^. Notably, the Wnt/β-catenin signaling pathway plays a particularly crucial role in these processes. In vitro models, activation of the Wnt pathway, such as with Wnt3a, during the pluripotent to mesoderm stage significantly increases cardiomyocyte differentiation rates in both EB and 2D methods^51,52^. Furthermore, inhibiting the Wnt pathway post-mesoderm formation enhances cardiac differentiation efficiency from pluripotent stem cells^53,54^. These insights provided the basis to formulate strategies of activating and inhibiting the Wnt pathway in cardiac model research, including advancements in self-organizing human heart models^7,8,13,55–57^.

FGFs can regulate both the pluripotency of stem cells and heart development through the modulation of specific gene expression profiles^15,16^. FGF2, for instance, when combined with BMP2, promotes cardiomyocyte differentiation from both ESCs and iPSCs ^58,59^. Additionally, FGF2 also contributes to the differentiation of stem cells into cardiac fibroblasts ^60^ and cardiac organoids^9,13^. Furthermore, FGF2, maintains pluripotency by modulating Wnt signaling through activation of the PI3-K/GSK3 pathway ^61,62^. Previous studies have demonstrated the cross-talk between FGF and Wnt pathways. For example, GSK-3β, which targets β-catenin for proteasome-mediated degradation, is also phosphorylated by AKT ^63^. Wnt signaling can confer competence for FGF signaling, while FGF signaling can also modulate Wnt signaling ^64–66^. The protocol described here leverages orchestrated cardiac signaling, including synergistic activation and inhibition of FGF and Wnt pathways, illustrating their intricate interactions during heart formation (Fig. 1 and 2). Our study emphasizes the significant role of these multiple cardiac signaling pathways, particularly how synergistic effects of FGF and Wnt activation and inhibition promote chamber formation and stability within CCOs (Fig. 1b, 1c and 2a). This orchestrated modulation not only enhances our understanding of cardiac developmental biology but also provides novel insights into the molecular mechanisms underlying organoid maturation and tissue morphogenesis. In another manuscript, we report the synergistic modulation of FGF and Wnt to generate a functional 3D proepicardial organoid, confirming the effectiveness and importance of this synergistic modulation approach. Our results may encourage future optimization for better cardiac organoids generation methods.

Our studies also have limitations worthy of future investigations. First, while we orchestrate multiple cardiac signals, including AA, RA, FGF, BMP and Wnt pathways, further detailed investigations are needed to understand the crosstalk between these pathways. Secondly, although our study shows that blocking FGFR leads to alterations in CCOs cell fate (Fig. 4e-4i), consistent with previous reporter^40^, the precise mechanisms driving these changes remain to be fully understood. Investigations into the molecular pathways and regulatory networks responsible for these alterations may generate much needed knowledge regarding human cardiac development. Lastly, our CCOs only resemble part of the heart chamber formation stage; for instance, the looping process is lacking.

In conclusion, our study offers a detailed understanding of the molecular mechanisms driving chamber formation in cardiac organoids from hPSCs. The CCOs generated through our optimized protocol hold significant potential for modeling drug screening, cardiac diseases, and regenerative medicine applications. Future research should focus on refining these models to further enhance their physiological relevance and utility in translational research.

## Supporting information

Bright-Field Microscopy Video of Live Stable Chambered Cardioids Beating on Day 60

Bright-Field Microscopy Video of Live Stable Chambered Cardioids Beating on Day 16.5

Live Calcium Imaging in Stable Chambered Cardioids on Day 50

## Acknowledgements

We thank all laboratory members for their help and discussions. We thank the Imaging Facility of Westlake University for assistance in organoid imaging. We thank the Flow Cytometry Facility of Westlake University for assistance in cryosection and live cell sorting. This work was supported by Westlake Laboratory (Key R&D Program of Zhejiang, 2024SSYS0032), and the National Natural Science Foundation of China (92068201). We also would like to thank all the Cell Fate Control Lab members for their support.

## Author contributions

D.P. provided financial support. X.Z. developed the method for generating chambered cardiac organoids (CCOs) and designed the experiments. D.P. and X.Z. conceived the project. X.Z. and D.P. made the formal analysis. H.Z. and T.K. analyzed the high throughput data set. F.W. performed the biochemical experiments and image processing. X.Z., F.W., and H.Z. wrote the manuscript draft. D.P. and X.Z. modified the draft. All authors contributed to the manuscript preparation.

## METHODS

### REAGENT or RESOURCE SOURCE IDENTIFIER

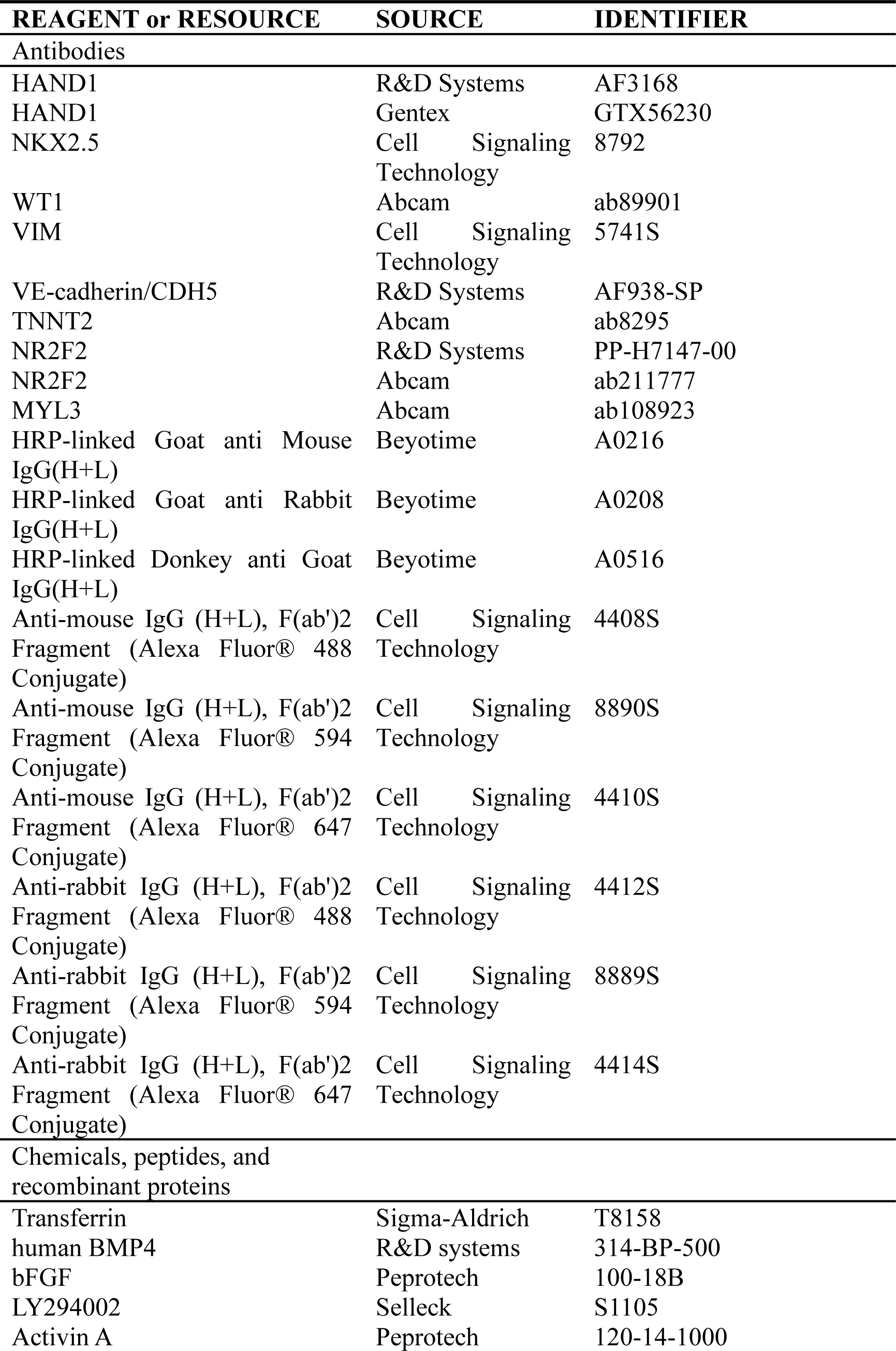
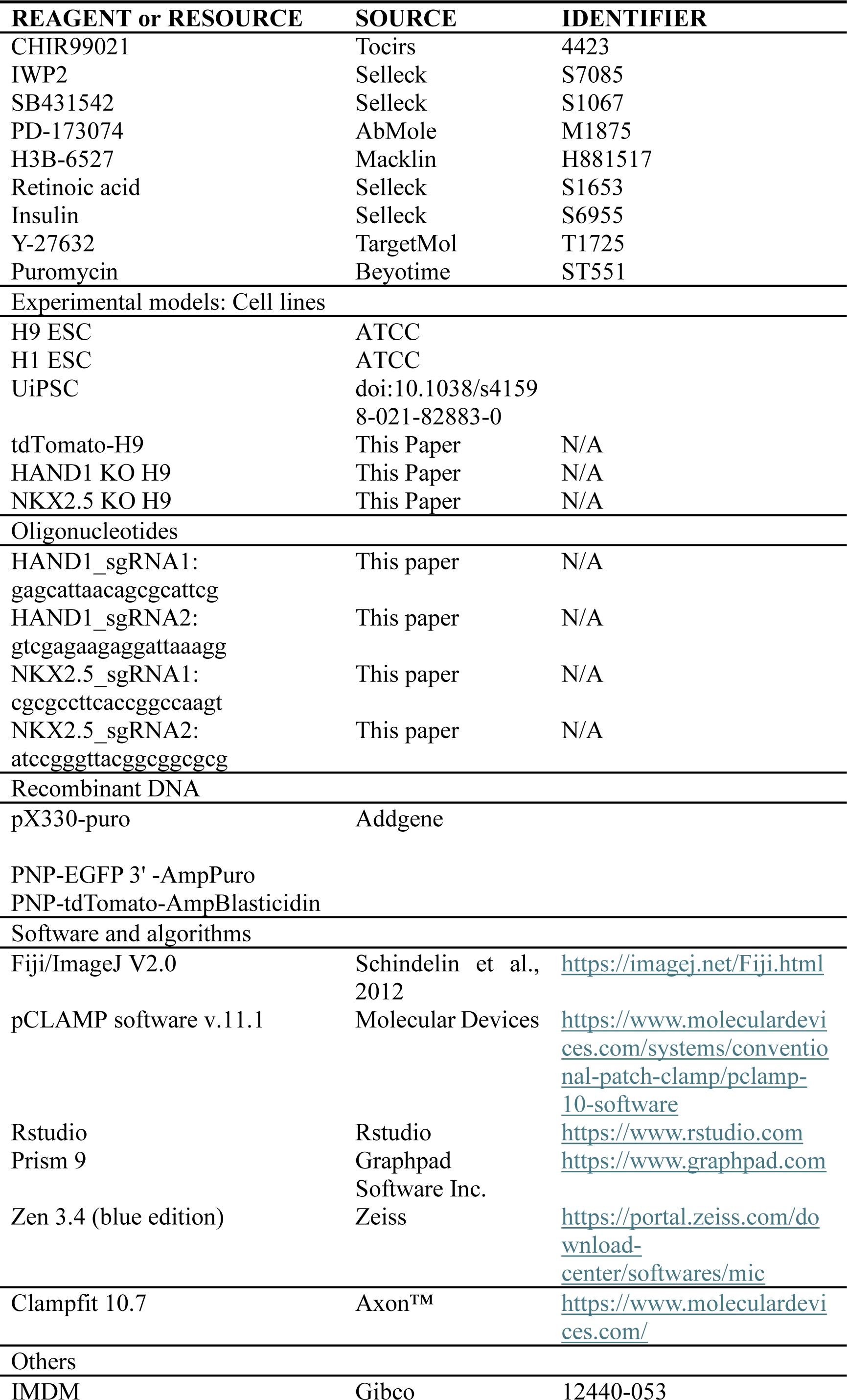
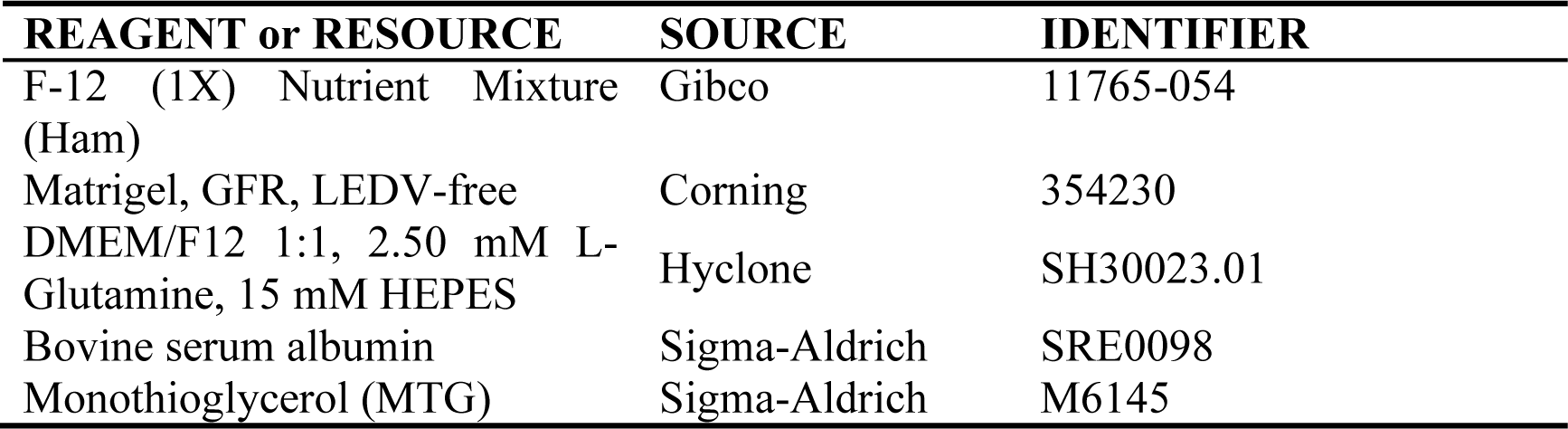

### Data and Code Availability

RNA sequencing data have been deposited in the NCBI Gene Expression Omnibus and are accessible through GEO accession numbers GSE269571 and GSE269572 (https://www.ncbi.nlm.nih.gov/geo/query/acc.cgi?acc=GSE269571, https://www.ncbi.nlm.nih.gov/geo/query/acc.cgi?acc=GSE269572).

## EXPERIMENTAL MODELS AND SUBJECT DETAILS

### Cell Lines

HEK293T cells (CRL-3216) and human ESCs (H9 and H1) were obtained from American Type Culture Collection (ATCC). HEK293T cells were cultured in Dulbecco’s modified Eagle’s medium (DMEM, Gibco) medium supplemented with 10% fetal bovine serum (FBS) and antibiotics. Human ESCs and home-made UiPSC^20,26^ were cultured in mTeSR1 medium (Stemcell Technologies, #85850) on Matrigel-coated (Corning, #354230) plates and passaged using either TrypLE Express Enzyme (Thermo Fisher, #12563029) or 0.5mM EDTA-DPBS (Sigma-Aldrich, #E6758) every 2∼4 days at 70%∼90% confluency. Cells were routinely tested for absence of Mycoplasma contaminations. All cell lines were cultured at 37 °C in a humidified atmosphere containing 5% CO_2_. In addition to specific mentions, all cell lines used were H9 cell lines.

### Generation of 2D cardiomyocytes and 3D chambered cardiac organoids

hPSCs were detached using Accutase (Stemcell Technologies, #07920) and resuspended in mTeSR1 medium supplemented with 10 μM ROCK inhibitor Y-27632. Cells were then seeded into Ultra-Low-Attachment U-bottom 96-well plates (Corning #7007 or Thermo Fisher #174929). For 2D differentiation, cells were seeded on a Matrigel-coated 24-well plate. The differentiation process involved a series of medium changes and specific supplements to promote cardiomyocyte and organoid formation. Detailed information about the medium composition and differentiation conditions is available in the protocol, which will be published separately.

### Generation of 2D epicardium

150,000 cells were seeded on a MG-coated 24 well. On day 0, medium was changed with CDM containing 30ng/mL FGF2, 3ng/mL BMP4, 10 μM CHIR99021 and 5 μM LY294002. After a 36h ∼ 40h nourishment, embryonic bodies were then incubated for 2 days with CDM medium containing 3ng/mL BMP4, 5μM IWP2 and 1μM BMS. From day 3.5 to day 6.5, spheroids were cultured in CDM medium containing 10 μg/mL insulin, 10ng/mL BMP4, 5μM CHIR 99021 and 1 μM Retinoic Acid. For maintaining epicardium, CDM medium containing 10 μg/mL insulin and 5μM SB-431542 were employed from day 6.5 onwards and refresh organoids every 2 days.

### Cryosectioning and immunostaining

Organoids were subjected into 30% sucrose in PB for dehydration and 4% PFA (Beyotime, #P0099) for fixation before section. For sectioning, organoids were embedded into O.C.T. (Sakura, #4583) and dissected into 10um slices by Leica cryostat at -22℃∼-20℃ for immunofluorescence.

Cryosections were post-fixed in 4% PFA and permeabilized in 0.2% Triton X-100 (Sigma-Aldrich, #T9284) in block solution (Beyotime, #P0102) for 15min each. After washing by DPBS, cryosections were blocked by using the above-mentioned blocking solution before incubated with the primary antibody diluted in primary antibody dilution medium (Beyotime, #P0103) for 4h at room temperature (RT) or overnight at 4℃ according to the instructions. Sections were then washed by PBS/0.1% Tween20 for three times and stained with the secondary antibody diluted in secondary antibody dilution medium (Beyotime, #P0108) for 2h at RT. Sections were prepared for observation by co-stained with DAPI (Abcam, #ab104139) for 5min at RT. Images were token from no less than 3 organoids by using inverted confocal microscope (Zeiss, LSM900 or LSM800).

### Generation of HAND1 and NKX2-5 knock out cell lines

CRISPR/Cas9 system was employed for knock out strategy. sgRNAs were identified using the website (http://crispor.tefor.net/crispor.py) and cloned into pX330-puro after hU6-sgScaffold site. Targeting efficiency of sgRNAs was validated in 293T cell line. Cells were transfected with 2μg pX330-sgRNAs and 4μg Donor per 1×106 cells using the Nucleofector™ 2b (Lonza-BioResearch, program B16). Post nucleofection, cells were incubated in mTeSR1 supplemented with 10 mM Y-27632 for 24h and then selected with 0.1ng/mL puromycin (Beyotime, #ST551) for 48h∼72h following with mTeSR1 plus 10 mM Y-27632. Once cell growth reached 50% confluency, ESCs were de-attached by Accutase and seeded into 96w-plate at 0.8cell per well. Single colonies will be macroscopic after 4∼5days and can be transferred after 10 days for the following genotyping using two different pairs of primers to be confirmed a successful knock out.

### Transmission Electron Microscope

Samples were fixed for 30min at room temperature or overnight at 4 °C in 150mM HEPES buffer (pH 7.2) containing 2.5% glutaraldehyde and 2% formaldehyde. Pre-fixed samples were washed using 0.1M PB/Cacodylate buffer (pH 7.2-7.4). After post-fixation in 1% osmium tetroxide for 1h at 4°C and in 1% uranyl acetate for 1h at room temperature or overnight at 4°C, samples were dehydrated in a graded series of ethanol (50%, 70%, 90%, 95%, and twice in 100%) and embedded in EPON12 resin at 60℃ for 24h to 48h. Sectioning was performed into 70um at a UC7 microtome (Leica) by using a diamond knife. After stained using basic dimethylamine blue and compound red, sections of 10 nm in thickness were collected on formvar-coated copper single-slot grids, post-stained with 2% uranyl acetate and lead citrate and observed in a 120 kV Transmission Electron Microscope (Thermo scientific, Talos120).

### Calcium imaging

Calcium flux was imaged by recording the fluorescence signal of real-time Fluo-4 AM. Briefly, after washing with DPBS, organoids were incubated with culture medium supplemented with 2 μM Fluo-4 AM (Beyotime, #S1060) and 0.1% Pluronic F-127 (Beyotime, #St501) for 30 minutes at 37 °C. The organoids were then transferred into a glass bottom plate (Cellvis, #P96-1.5H-N) with fresh medium for a 15-minute equilibration period at 37 °C. Organoids were prepared for recording by using a multi-mode Spinning Disk Confocal System (Olympus, Spin SR10) for more than 30s at a frame rate of 50ms per frame. Time-series images were analyzed using Fiji/ImageJ and processed using GraphPad Prism 9. The baseline fluorescence intensity (F) was calculated using asymmetric least squares smoothing. The change in fluorescence (ΔF/F) was determined by the formula

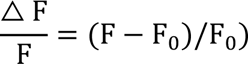

### Electrophysiology

The methodology was as previously described ^67^. Organoids were dissociated into single cells using the STEMdiff Cardiomyocyte Dissociation Kit (Stemcell Technologies, #05025). Action potentials (APs) were triggered at 1 Hz by 5 ms suprathreshold stimulations. Briefly, APs were recorded at room temperature (RT) using the whole-cell patch clamp technique by an Axopatch 700A amplifier and Digidata 1550B digitizer (Axon Instruments, Foster City, CA, USA). External solution contained the following: 132 mM NaCl, 4.8 mM KCl, 2 mM CaCl_2_, 1.2 mM MgCl_2_, 10 mM HEPES, and 5 mM glucose (pH was adjusted to 7.4 with NaOH). Internal solution contained the following: 110 mM KCl, 5 mM ATP-K_2_, 11 mM EGTA, 10 mM HEPES, 1 mM CaCl_2_, and 1 mM MgCl2 (pH was adjusted to 7.3 with KOH). Pipette series resistance was typically 1.5-3MΩ when filled with internal solution.

### RNA-seq and data analysis

Total RNA was extracted using the RNA-easy Isolation Reagent (Vazyme, #701) according to manufacturer instructions. Sequencing libraries were generated using NEBNext® UltraTM RNA Library Prep Kit for Illumina® and index codes were added to attribute sequences to each sample. Library quality was assessed on the Agilent Bioanalyzer 2100 system. The library preparations were sequenced on an Illumina NovaSeq platform (Illumina, USA). To analyze the gene expression, The raw sequencing data (Raw data) was filtered to get high quality sequencing data (Clean data). Use the Sequence Alignment Tool STAR to align with the human reference genome (reference genome source: https://ftp.ensembl.org/pub/release-109/fasta/homo_sapiens/dna/ and https://ftp.ensembl.org/pub/release-109/gtf/homo_sapiens/) to obtain the data that can be localised to the reference genome, and then use RSEM software to quantify the gene expression levels, and obtain the final normalisation matrix from the quantitative results. Then, DESeq2 (v. 1.26.0) was used for data normalization and differential expression analysis. Differentially expressed genes were defined by Wald test (Benjamini-Hochberg-corrected P-value < 0.05 and absolute fold change > = 1.5) and Likelihood ratio test (Benjamini-Hochberg-corrected P-value < 0.05) for time course experiments. Gene ontology analysis was performed using David (https://david.ncifcrf.gov/)

### PCA trajectory plots

Firstly, the gene expression matrix was analysed by PCA using the prcomp function in R language, and then the PCA scores were extracted. And according to the PCA analysis results of different samples at each time point, and connect the points of the same group through the trajectory line to indicate the trend of change between samples. Finally, PCA trajectory plots were drawn using the toolkit ggplot2 (v. 3.5.0).

### Heatmap

First, a DESeqDataSet object was created based on the gene expression matrix and experimental design information via the DESeqDataSetFromMatrix function in the R language package DESeq2 (v. 1.26.0), and then, the varianceStabilisingTransformation function was used to perform a variance stabilising transformation to obtain the processing data matrix. Finally, the heatmap function is used for plotting the heatmap.

### Single-cell (sc)RNA-seq and bioinformatic analysis

Organoids were dissociated into single cells using the STEMdiff Cardiomyocyte Dissociation Kit (Stemcell Technologies, #05025). The cell suspension was loaded into Chromium microfluidic chips with 3’ (v2 or v3, depends on project) chemistry and barcoded with a 10× Chromium Controller (10XGenomics). RNA from the barcoded cells was subsequently reverse-transcribed and sequencing libraries constructed with reagents from a Chromium Single Cell 3’ v2(v2 or v3, depends on project) reagent kit (10X Genomics) according to the manufacturer’s instructions. Sequencing was performed with Illumina (HiSeq 2000 or NovaSeq, depends on project) according to the manufacturer’s instructions (Illumina). The R language software package Seurat was used to process the scRNA data of the 2D group and 3D group, as well as the +PD group and -PD group. Low quality cells (< 300 genes/cell, > 9000 genes/cell, < 3 cells/gene and > 25% mitochondrion genes) were filtered out from the dataset. After quality control, an expression matrix containing 40,191 cells (2D and 3D) and 27,489 genes, and an expression matrix of 33,348 cells (+PD and -PD) and 27,583 genes were obtained, respectively.

Next, the data were normalized using the default parameters of the log normalization method, and the top 2000 highly variable genes were identified using the FindVariableFeatures function. The ScaleData function was used to Z-score transform gene expression. Dimensionality reduction analysis was then performed using the RunPCA function. Clustering was based on the 20 most important principal components (PCs). Batch problem removal was performed using the R package Harmony v0.1.0. Different resolutions were set, and clustree was used to observe the clustering effect. Finally, the resolutions were set to 0.9 and 0.7, respectively, and a unified manifold approximation and projection (UMAP) visualization image was constructed using the same number of PCs as the relevant clusters.

### Enrichment analysis of marker genes according to 2D cultures vs.3D cultures

We used FindMarkers function to find marker genes for cell types unique to the 3D cultures versus other cell types, and screened the significant marker genes according to Padj<0.05, |log2FC|>1. The screened genes were subjected to David(https://david.ncifcrf.gov/) enrichment analysis, and according to the enrichment results, the significant pathways with FDR less than 0.05 were screened. The analysis was then performed to reveal the potential biological functions of the 3D cultures.

### Quantification and statistical analysis

All analyses were performed using GraphPad software and all raw data was collected in Microsoft Excel. All data presented a normal distribution. Statistical significance was evaluated with a standard unpaired Student t-test (2-tailed; n.s. no significance, *p < 0.05, **p < 0.01, ***p < 0.001, ****p < 0.0001) when appropriate. For multiple-comparison analysis, one-way ANOVA with the Tukey’s or Dunnett’s post-test correction was applied when appropriate (n.s. no significance, *p < 0.05, **p < 0.01, ***p < 0.001, ****p < 0.0001). All data are presented as Mean ± SEM and represent a minimum of 3 independent experiments with at least 3 technical replicates unless otherwise stated. All micrograph images are representative of at least 6 independent experiments per condition/marker and calcium transient graphs are representative of 6 independent experiments.

