## Extended Data for "Self-Assembled Chambered Cardiac Organoids for Modeling Cardiac Chamber Formation and Cardiotoxicity Assessment"

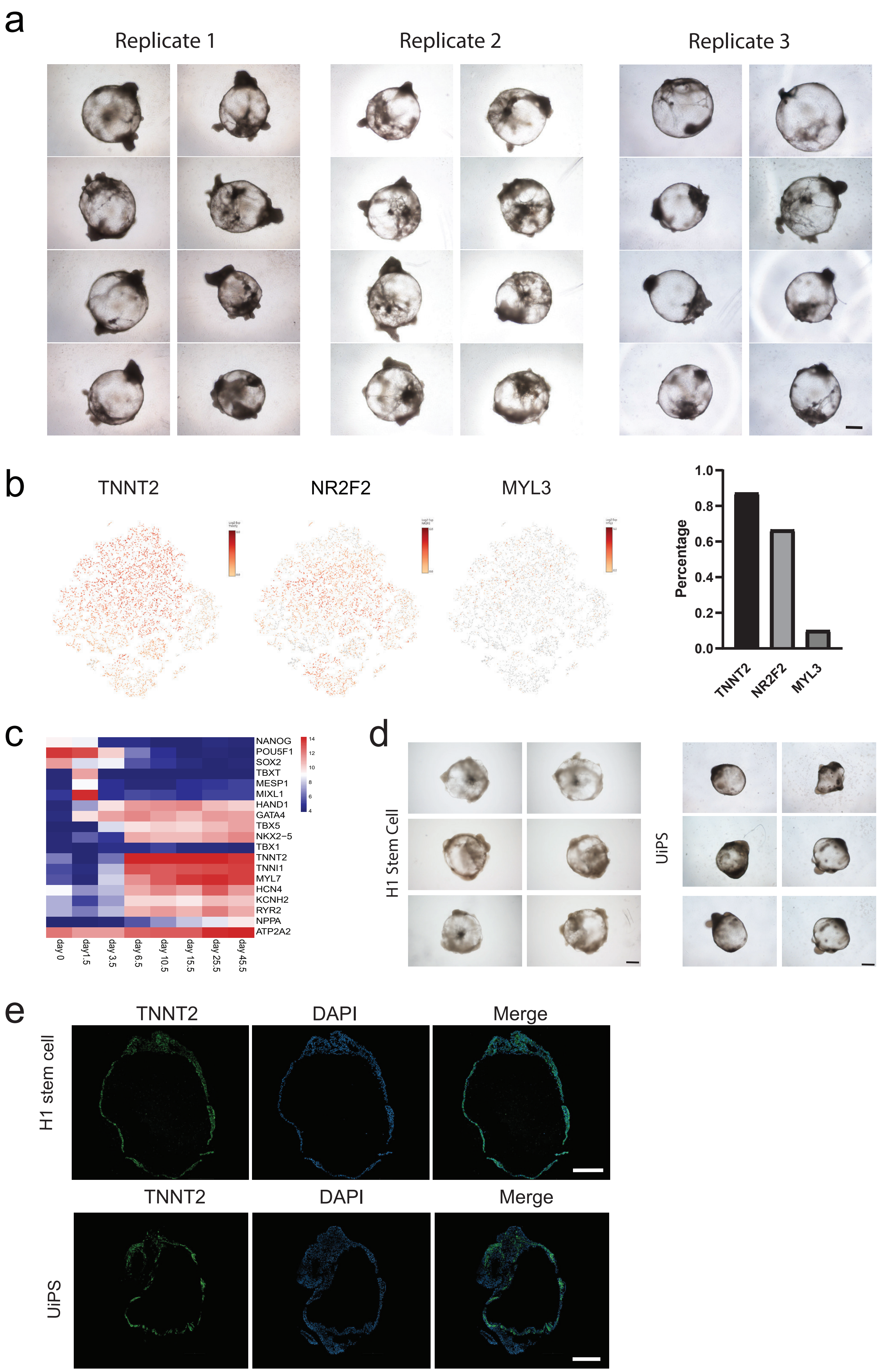
a
Replicate 1
Replicate 2
Replicate 3
b
TNNT2
NR2F2
MYL3
Percentage
1.0
0.8
0.6
0.4
0.2
0.0
TNNT2
NR2F2
MYL3
c
NANOG
POU5F1
SOX2
TBXT
MESP1
MIXL1
HAND1
GATA4
TBX5
NKX2−5
TBX1
TNNT2
TNNI1
MYL7
HCN4
KCNH2
RYR2
NPPA
ATP2A2
14
12
10
8
6
4
day 0
day1.5
day 3.5
day 6.5
day 10.5
day 15.5
day 25.5
day 45.5
d
H1 Stem Cell
UiPS
e
H1 stem cell
TNNT2
DAPI
Merge
UiPS

Extended Data Figure 1. Chamber formation of chambered cardiac organoids (CCOs) deprived from pluripotent stem cells, related to Figure 1.

a. Representative brightfield images from day 15.5, demonstrating the consistent emergence of cavity-containing structures across three separate biological replicates. Scale bar, 500 μm.
b. Cardiomyocyte, atrial, and ventricular cell proportions in CCOs, displayed in a UMAP plot.
c. Heatmap depicting the expression patterns of key genes during cardiomyocyte differentiation. VST, variance-stabilized transformed counts.
d. Representative brightfield images of CCOs derived from H1 and UiPS cell lines on day 15.5. UiPS, urine-induced pluripotent stem cells. Scale bar, 500 μm.
e. Cryosections of CCOs on day 16.5 showing the expression of the cardiomyocyte marker TNNT2 derived from H1 and UiPS cell lines. Scale bar, 200 μm.

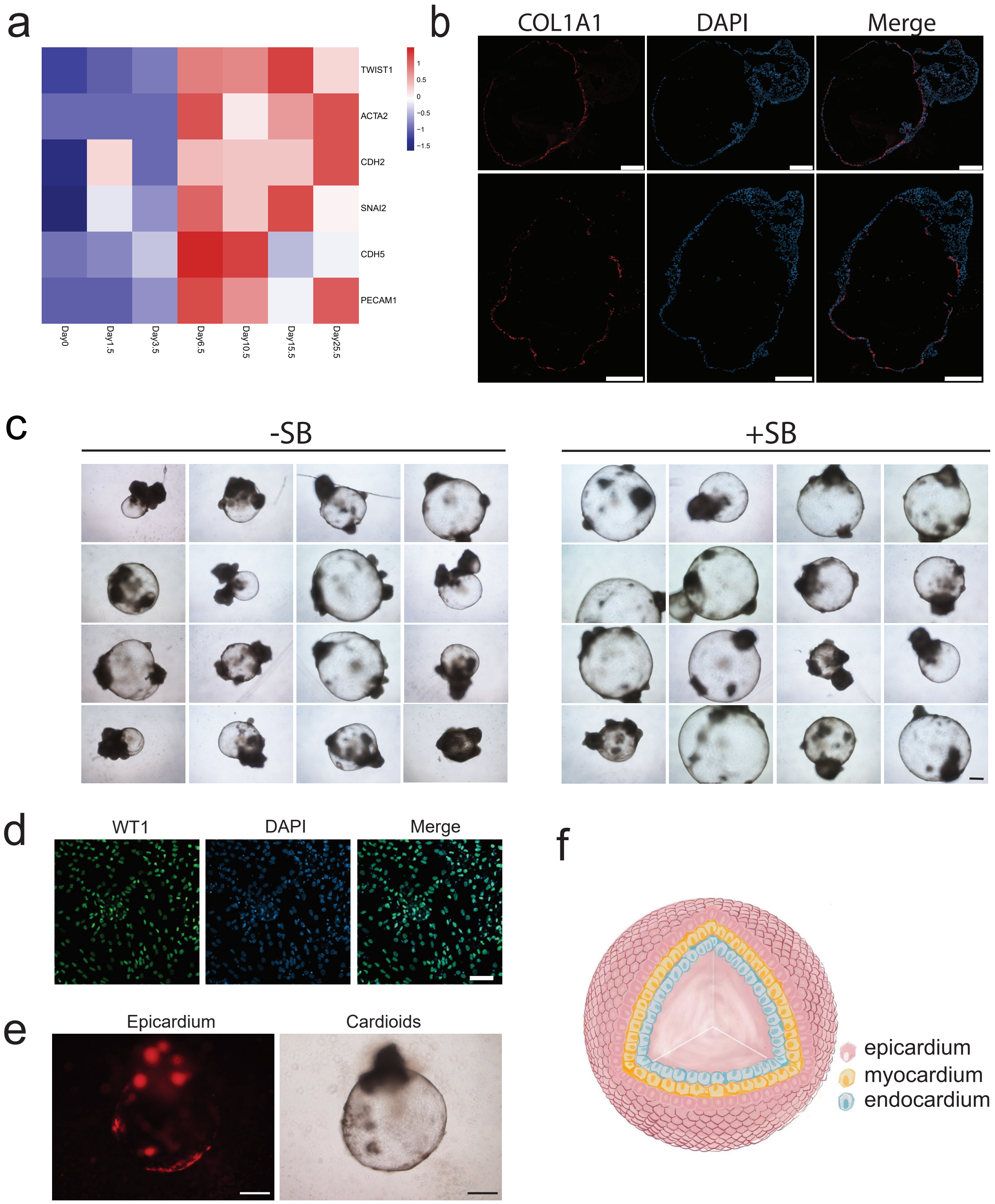
a
TWIST1
ACTA2
CDH2
SNAI2
CDH5
PECAM1
Day0
Day1.5
Day3.5
Day6.5
Day10.5
Day15.5
Day25.5
1
0.5
0
−0.5
−1
−1.5
b
COL1A1
DAPI
Merge
c
-SB
+SB
d
WT1
DAPI
Merge
e
Epicardium
Cardioids
f
epicardium
myocardium
endocardium

Extended Data Figure 2. CCO mirrors cardiac chamber formation stages, related to Figure 2.

a. Heatmap displaying gene expression of markers associated with endothelial-to-mesenchymal transition (EndoMT) during CCOs development from day 0 to day 25.
b. Cryosections of CCOs on day 38.5 showing the expression of the valve marker COL1A1. Scale bar, 200 μm.
c. Representative brightfield images of CCOs treated with or without 0.5 μM SB431542 for 20 days, captured on day 48.5.
d. Confocal images showing 2D epicardium formation with expression of the marker WT1 on day 9.5. Scale bar, 50 μm.
e. Live images of CCOs captured in tdTomato or brightfield. Scale bar, 500 μm.
f. A schematic diagram illustrating the three layers of CCOs: epicardium, myocardium, and endocardium.

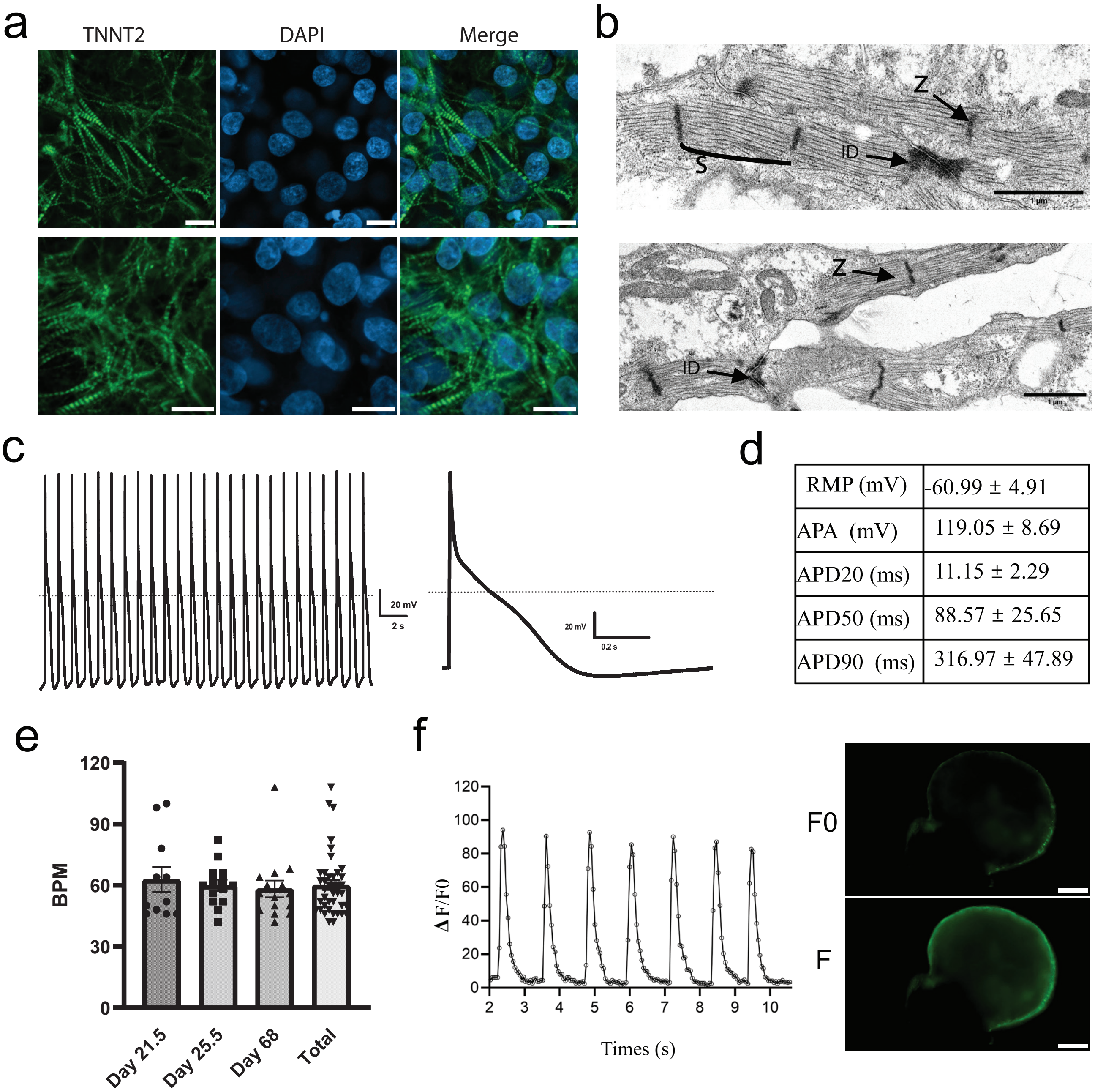


| | |
|---|---|
| RMP (mV) | -60.99 ± 4.91 |
| APA (mV) | 119.05 ± 8.69 |
| APD20 (ms) | 11.15 ± 2.29 |
| APD50 (ms) | 88.57 ± 25.65 |
| APD90 (ms) | 316.97 ± 47.89 |

Extended Data Figure 3. Functional characterization of CCOs.

a. Confocal images showing the sarcomeric organization within cardiomyocytes of CCOs on day 62. Scale bar: 10 μm.
b. Electron micrographs of CCOs on day 90.5, highlighting key structures: S, sarcomere; ID, intercalated disc; Z, Z-line.
c. Representative traces and average action potential recordings of CCOs on day 80.
d. Characteristics of action potentials observed in the data presented in (C). RMP, rest membrane potential; APD, action potential duration; APA, action potential amplitude.
e. Beating rate of cardiac organoids on different days. BMP, beats per minute. All bar graphs show mean ± SEM.
f. Calcium imaging analysis involved loading CCOs with Fluo-4-AM and monitoring fluorescence intensity over time. F/F0 represents fluorescence intensity relative to background levels. Scale bar, 500 μm.

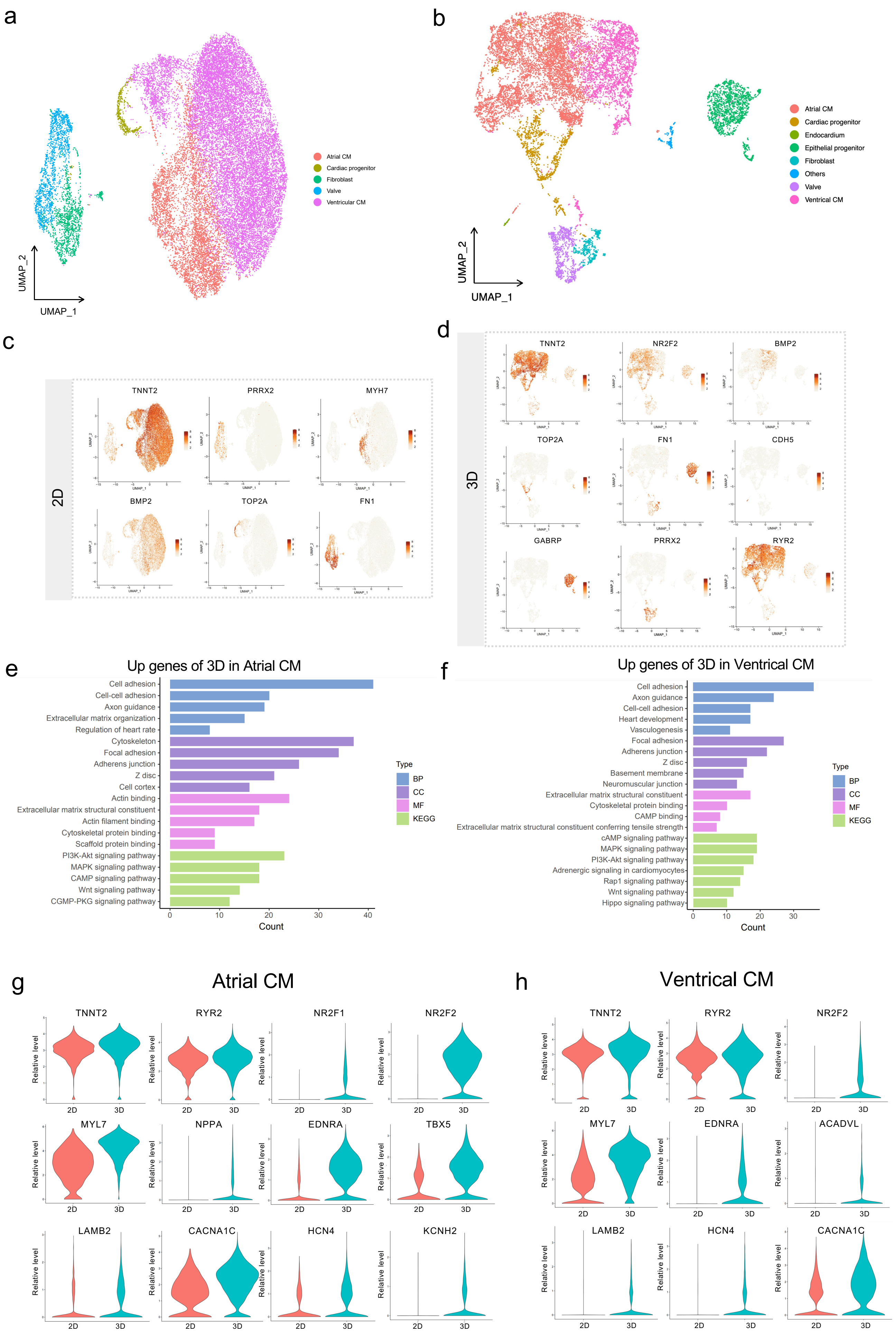
a
UMAP_2
UMAP_1
Atrial CM
Cardiac progenitor
Fibroblast
Valve
Ventricular CM
b
UMAP_2
UMAP_1
Atrial CM
Cardiac progenitor
Endocardium
Epithelial progenitor
Fibroblast
Others
Valve
Ventrical CM
c
2D
TNNT2
PRRX2
MYH7
BMP2
TOP2A
FN1
d
3D
TNNT2
NR2F2
BMP2
TOP2A
FN1
CDH5
GABRP
PRRX2
RYR2
e
Up genes of 3D in Atrial CM
Cell adhesion
Cell-cell adhesion
Axon guidance
Extracellular matrix organization
Regulation of heart rate
Cytoskeleton
Focal adhesion
Adherens junction
Z disc
Cell cortex
Actin binding
Extracellular matrix structural constituent
Actin filament binding
Cytoskeletal protein binding
Scaffold protein binding
PI3K-Akt signaling pathway
MAPK signaling pathway
CAMP signaling pathway
Wnt signaling pathway
CGMP-PKG signaling pathway
0
10
20
30
40
Count
Type
BP
CC
MF
KEGG
f
Up genes of 3D in Ventrical CM
Cell adhesion
Axon guidance
Cell-cell adhesion
Heart development
Vasculogenesis
Focal adhesion
Adherens junction
Z disc
Basement membrane
Neuromuscular junction
Extracellular matrix structural constituent
Cytoskeletal protein binding
CAMP binding
Extracellular matrix structural constituent conferring tensile strength
cAMP signaling pathway
MAPK signaling pathway
PI3K-Akt signaling pathway
Adrenergic signaling in cardiomyocytes
Rap1 signaling pathway
Wnt signaling pathway
Hippo signaling pathway
0
10
20
30
Count
Type
BP
CC
MF
KEGG
g
Atrial CM
TNNT2
RYR2
NR2F1
NR2F2
MYL7
NPPA
EDNRA
TBX5
LAMB2
CACNA1C
HCN4
KCNH2
Relative level
2D
3D
h
Ventrical CM
TNNT2
RYR2
NR2F2
MYL7
EDNRA
ACADVL
LAMB2
HCN4
CACNA1C
Relative level
2D
3D

Extended Data Figure 4. Comparative analysis of cellular diversity and gene expression profiles individually in 2D and 3D cell cultures, related to Figure 3.

a. UMAP visualization of the cell type annotation results of scRNA data for 2D cultures with colors representing cell types.
b. UMAP visualization of the cell type annotation results of scRNA data for 3D cultures with colors representing cell types.
c. UMAP scatter heatmap of marker genes for different cell types in 2D cultures with color shades representing expression.
d. UMAP scatter heatmap of marker genes for different cell types in 3D cultures with color shades representing expression.
e. Results of enrichment analysis of highly expressed genes in the 3D cultures compared to the 2D cultures in atrial cells.
f. Results of enrichment analysis of highly expressed genes in the 3D cultures over the 2D cultures in ventricular cells.
g. Comparison of the difference in expression of maturation marker genes in Atrial cardiomyocyte between the 2D cultures and 3D cultures, using violin plots to visualize the distribution of expression levels of these key genes.
h. Comparison of the expression differences of maturation marker genes in Ventricle cardiomyocyte between the 2D cultures and 3D cultures, using violin plots to visualize the distribution of expression levels of these key genes.

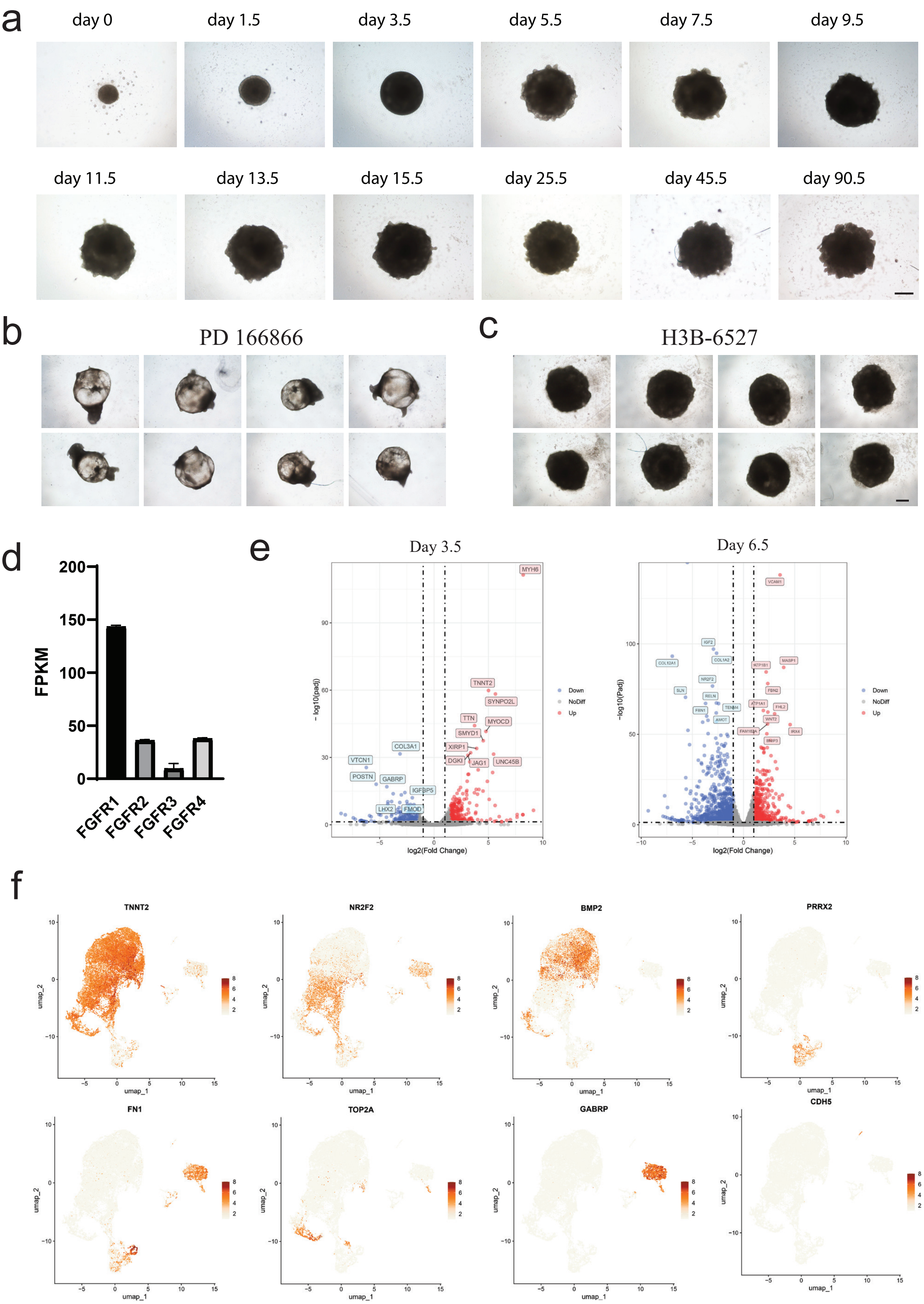
a
day 0
day 1.5
day 3.5
day 5.5
day 7.5
day 9.5
day 11.5
day 13.5
day 15.5
day 25.5
day 45.5
day 90.5
b
PD 166866
c
H3B-6527
d
FPKM
FGFR1
FGFR2
FGFR3
FGFR4
e
Day 3.5
Day 6.5
log2(Fold Change)
−log10(padj)
Down
NoDiff
Up
MYH6
TNNT2
SYNPO2L
MYOCD
TTN
SMYD1
XIRP1
DGKI
JAG1
UNC45B
COL3A1
VTCN1
POSTN
GABRP
IGFBP5
LHX2
FMOD
VCAM1
f
TNNT2
NR2F2
BMP2
PRRX2
FN1
TOP2A
GABRP
CDH5
umap_1
umap_2

Extended Data Figure 5. The role of FGF in the chamber formation of CCOs, related to Figure 4.

a. Time-lapse brightfield images illustrating the formation of cardiac organoids in the absence of the FGFR1 inhibitor PD17307. Scale bar, 500 μm.
b. Representative brightfield images of CCOs induced with PD166866 on day 17.5.
c. Representative brightfield images of cardiac organoids induced with H3B-6257 on day 17.5. Scale bar, 500 μm.
d. Expression levels of FGFR family genes on day 0. FPKM, Fragments Per Kilobase of transcript per Million mapped reads.
e. Volcano plot comparing gene expression profiles at the cardiac mesoderm stage on day 3.5 and the chamber formation stage on day 10.5, with PD treatment serving as the control.
f. Marker Gene Expression: A UMAP scatter heatmap illustrates the expression of marker genes for different cell types, with color gradients representing expression levels.

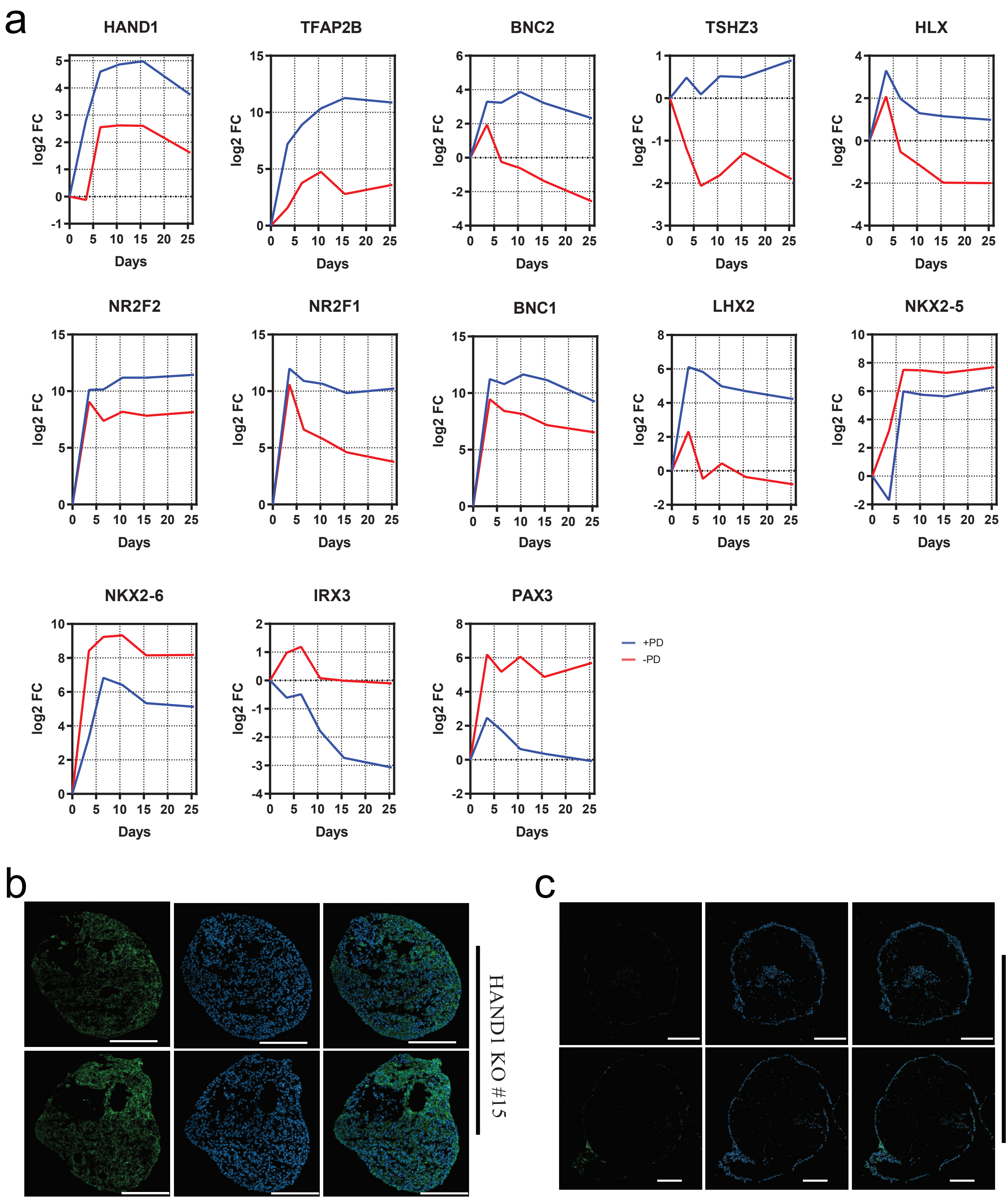
a
HAND1
TFAP2B
BNC2
TSHZ3
HLX
NR2F2
NR2F1
BNC1
LHX2
NKX2-5
NKX2-6
IRX3
PAX3
log2 FC
Days
+PD
-PD
b
HAND1 KO #15
c
NKX2-5 KO #24

Extended Data Figure 6. HAND1 is Crucial for the Formation of CCOs, related to Figure 5

a. Gene expression analysis of 13 selected genes (log2 fold-change vs. day 1.5) in cardiac organoids formation with or without PD173074. Blue line indicates with PD173074, while red line indicates without PD.
b. Confocal cryosection images of HAND1 knockout (KO) #15 cardiac organoids stained with TNNT2. Scale bar, 200 μm.
c. Confocal cryosection images of NKX2.5 knockout (KO) #24 cardiac organoids stained with TNNT2. Scale bar, 200 μm.

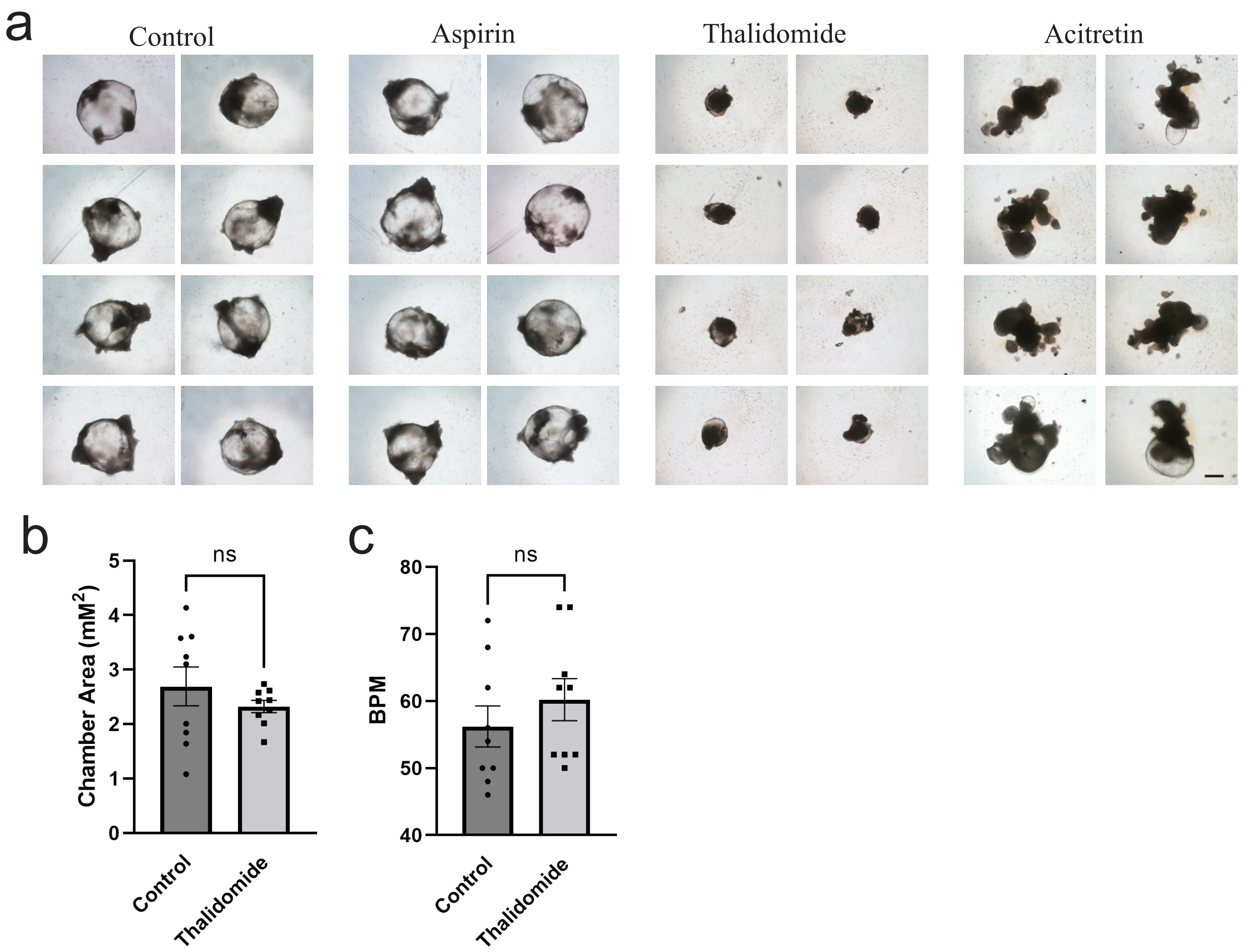
a
Control
Aspirin
Thalidomide
Acitretin
b
Chamber Area (mM2)
ns
0
1
2
3
4
5
Control
Thalidomide
c
BPM
ns
40
50
60
70
80
Control
Thalidomide

Extended Data Figure 7. Cardiotoxicity assessment platform using CCOs, related to Figure 7.

a. Representative brightfield images of CCOs treated with various drugs on day 20.5. Aspirin, 30 μM; thalidomide, 10 μg/ml; acitretin, 50 nM. Scale bar, 500 μm.
b. Quantification of the chamber areas of CCOs captured on day 36.5, treated with Thalidomide from day 15.5.
c. Beats per minute of CCOs on day 36.5, treated with Thalidomide from day 15.5.
